# Knockout and re-expression system for mutant analysis in primary mouse T cells

**DOI:** 10.64898/2026.08.25.746944

**Authors:** Vasileios Morfos, Meredith C. Frie, Daniel Peschkov, Juliana Wagner, Björn F. Lillemeier, Joanna Brzostek

## Abstract

We describe here an efficient method for gene editing in mouse T cells, based on well-established, high-efficiency retroviral transduction protocols. Our platform allows analysis of mutant phenotypes in primary murine T cells *in vitro* and *in vivo*. This approach uses a single retroviral vector to simultaneously knockout an endogenous gene and ectopically express its mutant version. This knockout/re-expression vector can be used as the only plasmid to transduce Cas9-expressing T cells, or used together with a Cas9 retroviral vector to transduce T cells from any mouse strain. We validated the system for analysis of murine T cells by targeting key molecules in proximal T cell signaling, i.e. CD3γ and Zap70. We obtain high knockout and re-expression efficiencies in both Cas9-expressing and non-Cas9 T cells. Knockout efficiencies can be further improved by gRNA multiplexing.

Endogenous proteins compete with their ectopically expressed mutants or tagged versions for cellular location, protein interactions and cellular functions. Here, we quantified the incorporation of CD3γ-GFP into surface T cell receptor (TCR) complexes. Our data shows that the knockout and re-expression platform improves integration of CD3γ-GFP into the TCR. Therefore, eliminating competition between endogenous and ectopic proteins benefits analyses of protein assemblies and signaling pathways in primary T cells. Furthermore, we validated advantages of our system for mutant analysis using wild-type and mutant Zap70s. Zap70 mutants deficient in TCR binding or kinase activity show their phenotypes only in the absence of endogenous protein, further validating our knockout/re-expression approach. Most importantly, this system can be used to generate gene-edited primary T cells for *in vivo* studies, such as the quantification of anti-tumor responses. Our knockout and re-expression platform provides a useful gene editing tool for primary T cells in fundamental research and immunotherapy development.

## Introduction

T cells protect against pathogens and tumors. They kill infected or malignant cells, and produce cytokines that orchestrate systemic immune responses. Primary mouse T cells are irreplaceable for the study of T cell signaling and function. Murine T cells can be genetically engineered *ex vivo*, followed by comprehensive analyses *in vitro* or *in vivo*. The functions of genes of interests can be investigated using ectopic overexpression, gene knockouts or knockins. Loss of function analyses using knockouts have been instrumental for studying T cell biology, and CRISPR/Cas9 knockout protocols for primary mouse T cells are well-established^1–3^. However, the discovery of molecular mechanisms and precise mapping of sequence-function relationships require mutant analyses either by overexpression or gene knockin. Retroviral transduction is widely used for protein overexpression in primary T cells, with 60-70% transduction efficiencies reported^4,5^. However, the use of overexpression to study T cells is limited to determining protein localizations^6,7^, or functional analysis of dominant negative mutants^8^. Knockins provide a powerful approach for mutant analysis, but remain inefficient and technically challenging for primary mouse T cells.

CRISPR/Cas9 is the most common approach to edit genes in primary T cells^9^. In brief, Cas9 nuclease in complex with a single guide RNA (sgRNA) complementary to the targeted gene sequence binds and cleaves genomic DNA. The resulting double strand breaks are repaired by error-prone non-homologous end joining (NHEJ). NHEJ generates insertions or deletions that disrupt the coding sequence of the gene, resulting in a knockout. In the presence of homology repair DNA templates, Cas9-mediated double stranded DNA breaks can be repaired though homology-directed repair (HDR). Desired gene mutations are introduced into repair templates, resulting in a knockin. Several protocols have been established for efficient CRISPR/Cas9-mediated gene knockout in primary mouse T cells. Retroviral transduction can deliver sgRNA into T cells from Cas9 transgenic mice with more than 90% knockout efficiency^1^. However, use of a single retroviral vector containing Cas9 and sgRNA was reported to result in significantly lower knockout efficiencies^1^. This is likely due to the large size of Cas9 reducing transduction efficiency. Alternatively, electroporation can deliver Cas9 protein in complex with sgRNA (ribonucleoprotein complex, RNP)^10–12^ or deliver plasmids encoding Cas9 and sgRNA^13^. Both of these electroporation-based delivery methods result in high (80-90%) knockout efficiencies in mouse T cells.

Mutant knockins for primary mouse T cells analyses remain challenging. HDR is inherently less efficient than NHEJ, and low knock-in efficiencies of 0.5 - 20% have been reported for Cas9-based knockins^14^. Moreover, knockin efficiencies in primary mouse T cells are limited by toxicity of homology repair templates. Specifically, electroporation with single- or double stranded DNA repair template results in high cell death and low (<10%) editing efficiency^11,13^. Multiple strategies have been developed to enhance knockin efficiency, including Cas9 engineering, repair template optimization and improved delivery methods^15^. Recent improvements in adeno-associated virus (AAV) gene delivery significantly improved knockin efficiency in primary mouse T cells^16,17^. However, in contrast to retroviruses, AAVs are not routinely used to transduce murine T cells, and infection protocols are not available in most laboratories. In addition, the utility of AAV is limited by its low packaging capacity^18^ and immunogenicity^19^.

We have developed and validated a reliable and easy to implement method for mutant analysis in mouse T cells, based on well-established, high-efficiency retroviral transduction protocols. Our approach uses a single viral vector to simultaneously knockout an endogenous gene and ectopically express its mutant version. We show that this single knockout and re-expression vector can be used alone to transduce Cas9 transgenic T cells, or any T cells when used together with a second retroviral vector encoding Cas9. For proof-of-principle, the system was used to study the T cell receptor (i.e. the TCR subunit CD3γ), and its associated kinase ZAP70. We observed high knockout and re-expression efficiencies, and specifically analyzed the effects of Zap70 signaling mutants *in vitro* and *in vivo*.

## Materials and Methods

### Cell lines

293T cells were maintained in DMEM with 10% COSMIC Calf serum (HyClone, Cytiva) and 100 U/ml penicillin-streptomycin, hence referred to as complete DMEM. MC38-OVA and MC38-null cells were maintained in complete DMEM with 400 μg/ml G418. EL4 cells were maintained in RPMI1640 with 5% COSMIC Calf serum and 100 U/ml penicillin-streptomycin.

### Mice

OT-I (C57BL/6-Tg(TcraTcrb)1100Mjb/J; Jax strain 003831), Cas9 (B6J.129(Cg)-Igs2tm1.1(CAG-cas9*)Mmw/J; Jax strain 028239) and Rag1KO (B6;129S7-Rag1tm1Mom/J, Jax strain 002096) mice were obtained from The Jackson Laboratory. OT-I Cas9 mice were bred by crossing OT-I with Cas9 mice. Mice were maintained under specific pathogen-free conditions in vivariums at the Salk Institute or University of Freiburg. Experiments were performed with 6- to 12-week-old male and female mice.

### Molecular biology

Standard molecular biology techniques were used to generated the vectors, based on pSIR-GFP^20^, pSpCas9(BB)-2A-GFP^21^ and MSCV.

pSIR-GFP was a gift from Hodaka Fujii (Addgene plasmid #51134), pSpCas9(BB)-2A-GFP (PX458) was a gift from Feng Zhang (Addgene plasmid # 48138).

sgRNAs targeting murine CD3γ, CD3d and Zap70 were designed using CHOPCHOP^22^. gRNA sequences are listed below:

Mock gRNA: AAAAAGTCCGCGATTACGTC

CD3γ gRNA_1: AAGAGACTGACATGGAGCAG

CD3γ gRNA_2: CTGGTGATCTCTCTTCTTCA

CD3δ gRNA: GAGTCCAGGAGCACATACTT

Zap70 gRNA: TGCTCTAGTGCATCGCCCTG

### Retroviral Transduction of Mouse OT-I Cas9 or OT-I Cells

ScreenFect A-plus (ScreenFect GmbH, S-6001) was used to transfect 293T cells with pCL-Eco and retroviral transfer plasmids for virus production. The following protocol is for transfection of one well in a 6-well plate to generate supernatant for transduction of 2×10^6^ T cells per well in 24 well plate; the volumes, DNA amounts and cell numbers can be scaled as required. 1.5 μg transfer plasmid and 0.5 μg pCL-Eco was added to 120 μl ScreenFect Dilution Buffer; 4 μl ScreenFect A-plus was added to 120 μl ScreenFect Dilution Buffer. The diluted ScreenFect A-plus was added to the diluted DNA and immediately mixed using 10 rapid pipette strokes, followed by 20 min incubation at room temperature (RT). During the 20 min incubation, 293T cells were detached using trypsin-EDTA. The trypsin was quenched by adding complete DMEM. 29T3 cells were the counted and cell concentration was adjusted to 1×10^6^ cells/ml using complete DMEM. 1.25ml (1.25×10^6^ cells) of cell suspension was added to the 240 μl transfection complexes, mixed by pipetting and transferred to TC-treated 6 well plate. Next day, 0.5 ml complete DMEM was added to the transfected 293T cells.

RPMI 1640 with 10% fetal calf serum, 50μM β-mercaptoethanol, 1× non-essential amino acids, 1× sodium pyruvate, 1mM HEPES, and 100U/ml penicillin-streptomycin, hence referred as complete RPMI, was used for culture of primary mouse T cells. Splenocytes and lymph node cells were isolated from OT-I Cas9 or OT-I mice, and dissociated into cell suspension using a 1-ml syringe plunger and a 70-μm cell strainer. Red blood cell lysis was performed by 5 min incubation with ACK buffer, followed by quenching using 10 ml complete RPMI and centrifugation. Lymphocytes and splenocytes from one mouse were incubated in 12 ml complete RPMI with 0.5 μM SIINFEKL peptide. After 24 h, density gradient centrifugation was performed using Pancoll mouse (PAN Biotech, P04-64100), and cells were transferred into 14 ml fresh complete RPMI with 50 U/ml recombinant human IL-2 (rhIL-2).

The transduction was performed 36h after start of T cell stimulation, and approx. 48h after transfection of 293T cells. Polybrene (Sigma-Aldrich) was added to supernatants harvested from transfected 293T cells at 10 μg/ml final concentration, followed by 20min incubation at RT. 2×10^6^ T cells in 100 μl complete RPMI were added to supernatant, and transferred to one well in 24 well plate. The spin infection was performed at 1,250×g for 90 minutes at 32°C. After centrifugation, the supernatant was carefully removed and replaced with 1 ml of conditioned media with 50 U/ml rhIL-2. The conditioned media was prepared by mixing equal volumes of fresh complete RPMI medium and the filtered (0.45μm filter) supernatant from T cell culture.

One day after transduction, 1 ml fresh complete RPMI with 50 U/ml rhIL-2 was added to the T cells. Two days after transduction, 1 ml fresh complete RPMI with 50 U/ml rhIL-2 was added to the T cells, and puromycin (5 μg/ml) and/or blasticidin (25 μg/ml) was added. Four or five days after transduction, density gradient centrifugation was performed using Pancoll mouse, and cells were re-suspended in 2 ml complete RPMI with 50 U/ml rhIL-2. Cells were used for analysis approx. 24h after density gradient centrifugation.

### Antibody staining for flow cytometry analysis

For cell-surface antibody staining, antibodies were diluted in FACS wash buffer (FWB; 0.5% bovine serum albumin in phosphate-buffered saline (PBS)), and FWB was used for all the subsequent washing and re-suspension steps. Live/dead labeling was performed before the antibody staining, using a LIVE/DEAD Fixable Near-IR Dead Cell Stain kit (Invitrogen; L10119) diluted 1:1,000 in PBS for 10 min on ice, followed by FWB wash. The cell surface staining was performed on ice for 30 min. The following antibodies were used from cell-surface staining: PE or BUV395 conjugated anti-mouse CD8α (clone 53-6.7, BD Biosciences or Biolegend), PE conjugated anti-mouse TCRβ (clone H57-597, Biolegend), APC conjugated anti-mouse CD25 (clone PC61, Biolegend).

For intracellular staining to detect cytokines, total Zap70 or HA-tagged Zap70, samples were fixed in 4% paraformaldehyde for 20 min at RT after the live/dead and cell surface staining was performed as described above. The samples were then washed in 1× permeabilization buffer (eBioscience Permeabilization Buffer, Thermo 00-8333-56), incubated for 30 min at RT with antibodies diluted in 1× permeabilization buffer, followed by a wash in 1× permeabilization buffer, and a final wash and re-suspension for analysis in FWB. The following antibodies were used for intracellular staining with this protocol: APC or PE conjugated anti-mouse IFNγ (clone XMG1.2, Biolegend), PE-Cy7 conjugated anti-mouse TNFα (clone MP6-XT22, Biolegend), PE conjugated anti-human/mouse Zap70 (clone IE7.2, Invitrogen), Alexa Fluor 647 conjugated anti-HA.11 (clone 16B12, Biolegend).

For pErk staining, samples were fixed by adding an equal volume of 8% paraformaldehyde at the end of the stimulation. Cells were fixed for 20 min at room temperature, followed by centrifugation. The samples were then permeabilized in 0.2 ml ice-cold 90% methanol/PBS for 30 min, followed by two washes in PBS. Samples were incubated for 30 min at RT with antibody diluted in FWB, followed by a final wash and re-suspension for analysis in FWB. Alexa Fluor 647 conjugated anti-ERK1/2 phospho-Thr202/Tyr204 (clone 4B11B69, Biolegend) was used.

Flow cytometry analysis was performed on Aurora spectral flow cytometer (CyTek), with SpectroFlow used for acquisition and FlowJo versions 9 and 10 used for analysis.

### T cell activation assays using target cells

T cell activation assays using target cells were performed in U-bottom 96 well plate in 200 μl total volume of cRPMI at 37°C, 5% C0_2_. Target cells were labelled using 0.5 μM CellTrace Violet for 10 min. In experiments using EL4 cells, the target cells were pulsed for 1 hour using the indicated peptide concentrations at 37°C, 5% C02, followed by three washes. In experiments using M38 cells, the target cells were detached using trypsin-EDTA prior to the co-culture.

For assays measuring cytokine production and degranulation, 1×10^5^ transduced T cells (day 7 or 8) were co-cultured with target cells at 1:1 ratio. Co-culture was performed in the presence of PE-conjugated anti-mouse CD107a (clone 1D4B, Biolegend) antibody and 1× Brefeldin A (eBioscience Brefeldin A Solution, 00-4506-51, Invitrogen). For assays measuring target cell killing and CD25 upregulation, 1×10^5^ target cells were incubated with T cells at the indicated E:T ratios for 4h (EL4 cells) or 24h (MC38 cells).

### T cell stimulation using anti-CD3ε antibody

T cell activation assay using anti-CD3ε antibody was performed in U-bottom 96 well plate. Per well, 0.5×10^6^ T cells were resuspended in 100 μl RPMI without any additives (plain RPMI), and serum-starved for 1h at 37°C. Plates were then transferred to ice, and 20 μl of anti-CD3ε antibody (clone 145-2C11, Biolegend) diluted in plain RPMI was added to a final concentration of 4 μg/ml. Samples were incubated on ice for 10 min. 20 μl of anti-Armenian and Syrian Hamster IgG1 antibody (clone G94-56, BD Biosciences) diluted in plain RPMI was added to each well for a final concentration of 2 μg/ml, followed by 15 min incubation on ice. T cell stimulation was initiated by transferring the samples to 37°C water bath, and stopped by adding an equal volume of 8% paraformaldehyde.

### Microscopy

For H-2K^b^ surfaces, eight-well glass chambers (Lab-Tek II chambered coverglass #1.5 borosilicate, Nalgen Nunc International) were cleaned with plasma (Harrick Plasma) and sequentially coated with 0.1 mg/ml biotinylated poly-l-lysine, 20 μg/ml streptavidin, and 100 nM biotinylated H2-K^b^ OVA monomer (NIH tetramer facility). For antibody surfaces, eight-well glass chambers were coated with 0.5 mg/ml poly-l-lysine, dried overnight, and then incubated with 10 μg/ml anti-mouse CD3ε Antibody (clone 145-2C11, Biolegend) in HEPES-buffered saline (HBS: 20 mM HEPES pH7.4 plus 150 mM NaCl) for 2h at 37°C. Both coated glass chambers were kept in HBS prior to imaging. For TCR staining, cells were spun at 250 g, resuspended in Imaging Media (Hanks’ Balanced Salt Solution (HBSS) with 0.5 mM CaCl_2_, 2 mM MgCl_2_, 1% FBS), and stained with 17 µg/ml H57 Fab AF647 for 30 min on ice. Cells were washed with Imaging Media and spun at 250 g. Cells were transferred in Imaging Media and were kept on ice until imaging.

Cells were added to coated glass chambers containing Imaging Media equilibrated to 37°C. Cells were maintained at 37°C during image acquisition with a Life Imaging Services heating system. Imaging was performed on a Nikon Eclipse Ti2 inverted microscope platform with a 60x objective (CFI Apo TIRF oil immersion NA 1.49, Nikon) and a Kinetix 22 camera (Teledyne Photometrics). Image acquisition was performed with VisiView software (Visitron), and Images were edited with Fiji.

### In vivo tumor growth

MC38-OVA cells were detached with trypsin/EDTA, washed twice in ice-cold HBSS, and re-suspended in ice-cold HBSS at 2.5×10^6^ cells/ml. 100 μl of the cell suspension was injected subcutaneously into the right flank of Rag1KO recipients. On day 7, 2×10⁶ T cells were injected intravenously via the retro-orbital route. Tumors were measured every two days or daily using calipers. Tumor volume was calculated using the formula V = L x W^2^ where L is the longest tumor diameter and W is the shortest diameter. The experiments were performed in accordance with procedures approved by the Institutional Animal Care and Use Committee at Salk Institute for Biological Studies.

### Quantification and statistical analysis

Figure legends provide information statistical parameters such as the definition and value of n, and statistical significance. Differences were considered significant when p values were <0.05. Statistical significance in each figure was calculated as indicated in the figure legend. Only statistically significant differences are shown. Statistical analysis was performed in GraphPad PRISM 8.4.2

## Results

### Single vector knockout and re-expression platform for gene editing in mouse T cells

Analyses of mutant proteins frequently require knockout of the endogenous gene, as the presence of wild-type (WT) protein can mask mutant phenotypes. We developed a single retroviral vector system to simultaneously knockout the endogenous gene and ectopically express a mutant version. This strategy allows efficient generation of primary mouse T cells expressing mutant proteins in the absence of the endogenous WT proteins. For proof-of-principle experiments, we applied this strategy to the CD3γ subunit of the TCR complex, using T cells from OT-I TCR and Cas9 transgenic mice (hence referred to as OT-I Cas9). The T cell receptor (TCR) is a multimeric complex made of the ligand-binding TCRαβ dimer, and signal-transducing CD3ζζ, CD3ɛγ and CD3ɛδ dimers. The CD3 proteins contain cytoplasmic immunoreceptor tyrosine-based activation motifs (ITAMs), which become phosphorylated by the kinase Lck upon TCR ligation^23^. Phosphorylated ITAMs bind to and activate the kinase Zap70, which can then phosphorylate its downstream substrates leading to T cell activation^24,25^.

We designed a panel of pSIR-based retroviral vectors^20^ for simultaneous gene knockout and ectopic expression of proteins (Figure 1a). Each vector contains a murine U6 promoter driving transcription of either a control (mock) or a single guide RNA (sgRNA) targeting genomic *CD3γ* (CD3g_1 or CD3g_2). In addition, we designed a tandem sgRNA cassette (tandem), where the CD3g_1 and CD3g_2 sgRNAs are linked using the rice glycine tRNA sequence. After transcription, the tRNA is processed to release individual sgRNAs, allowing expression of two different sgRNAs from a single U6 promoter^26,27^. A PGK promoter drives expression of either GFP or a CD3γ-GFP fusion protein, and a puromycin resistance gene for antibiotic selection. Constructs expressing GFP contain the resistance gene downstream of an internal ribosome entry site (IRES), while constructs expressing CD3γ-GFP fusion protein use a T2A self-cleaving peptide^28^. The latter is shorter and keeps inserts sizes at similar lengths for better comparison.

**Figure 1.**
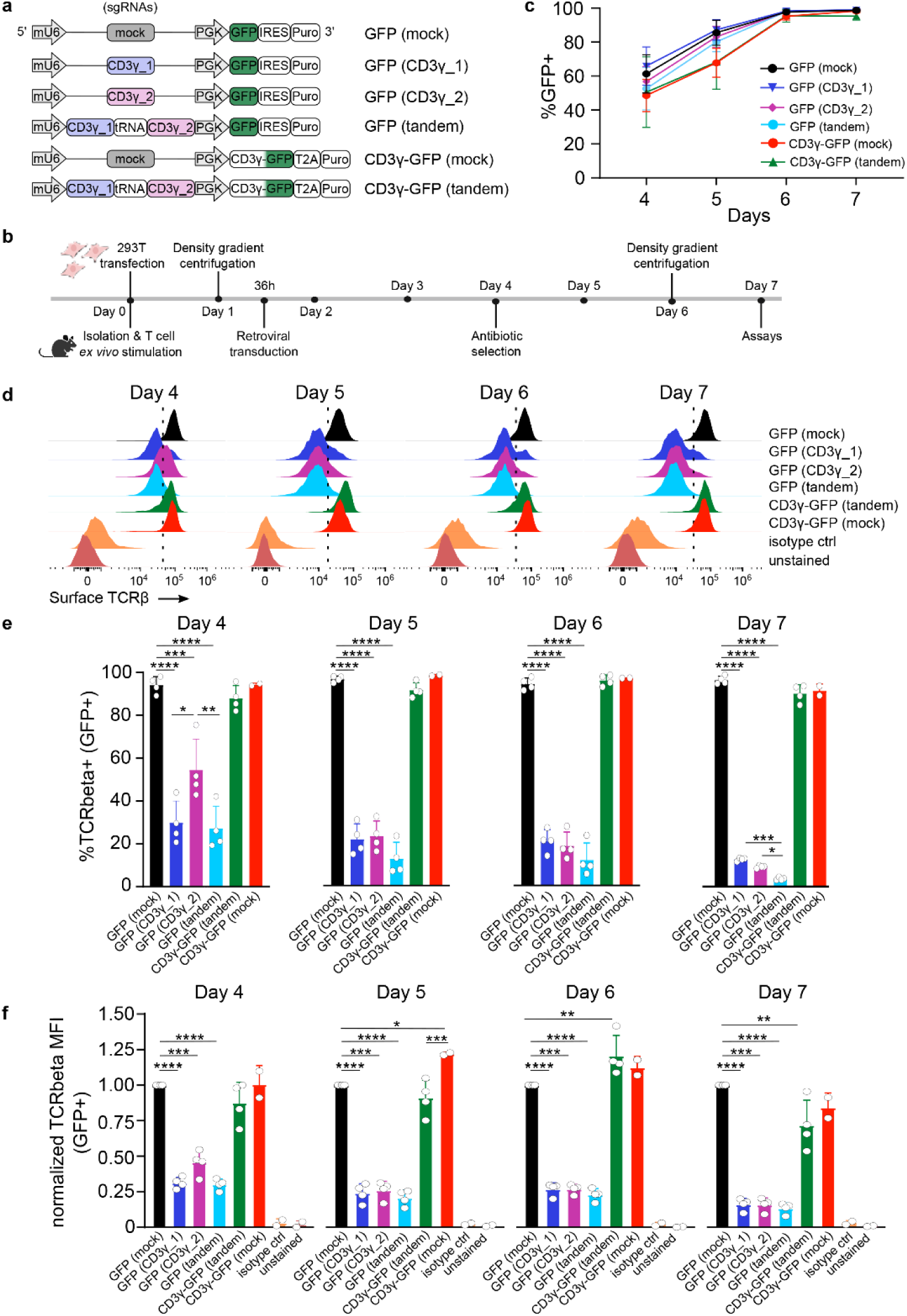
Single vector system to knockout and re-express CD3γ in mouse T cells. (**a**) Schematic overview of the retroviral vectors. (**b**) Timeline of retroviral OT-I Cas9 T cells transduction. (**c**) Percentage of transduced (GFP+) T cells over time. (**d**) Representative flow cytometry histograms showing surface TCRβ staining in GFP+ cells over time. (**e**) Percentage of TCRβ+ T cells within the GFP+ population over time. (**f**) TCRβ MFI on GFP+ population over time, normalized to GFP(mock) control. Data from two to four independent experiments, with one donor mouse per experiment. Data are presented as mean ± SD; in (e) and (f) each dot represents one independent experiment. Statistical analysis was performed using one-way ANOVA with Tukey’s multiple comparisons test.; P < 0.05 (*), P < 0.01 (**), P < 0.001 (***), P < 0.0001 (****).

We first tested the sgRNA knockout efficiency by analyzing the TCR surface expression. CD3γ is essential for the assembly and trafficking of the TCR/CD3 complex to the cell surface, and loss of CD3*γ* results in severe reduction of surface TCR^29^. The experimental timelines used here and in subsequent experiments are shown in Figure 1b. Briefly, OT-I Cas9 T cells were isolated and stimulated with antigenic SIINFEKL (Ovalbumin amino acids 257-267, hence referred to as OVA) peptide for 24h. Retroviral infection was performed 36-38h after T cell isolation. Antibiotic selection for transduced cells was started on day 4 after T cell isolation. GFP fluorescence and TCR surface expression were analyzed from day 4 to day 7, as indicators of transduction and knockout efficiencies, respectively. We observed 45-65% GFP+ cells on day 4, increasing to above 95% on day 7 (Figure 1c), indicating good transduction efficiency and effective antibiotic selection of transduced T cells. CD8 T cell numbers increased approximately 3-fold within four days after transduction (day 7). Similar numbers were obtained for all subsequent experiments using single vector transfections. The two single sgRNA constructs targeting *CD3γ*, as well as the tandem construct, induced robust loss of surface TCR, whereas mock sgRNA had no effect (Figure 1d and 1e, Supplementary Figure 1). TCR loss was apparent on day 4, and the proportion of TCR+ cells decreased over the next three days. The tandem construct had higher knockout efficiency than the individual gRNAs, resulting in TCR+ population at the detection limit on day 7 (Figure 1d and 1e). The tandem construct was therefore used in all subsequent experiments. TCR surface levels in single or tandem gRNAs transduced T cells were reduced more than 10-fold, but remained above unstained or isotype-stained T cells on day 7 (Figure 1f). This could be due to incomplete degradation of endogenous CD3γ or presence of non-canonical CD3γ-deficient TCR complexes^30^.

T cells were transduced with constructs encoding the tandem sgRNA and CD3γ-GFP to test if TCR surface expression can be restored. Re-expression of CD3γ-GFP resulted in >90% TCR positive cells at all time points, comparable to that of GFP(mock) or CD3γ-GFP(mock) controls (Figure 1e). Moreover, the TCR surface levels were similar for GFP(mock), CD3γ-GFP(mock) and CD3γ-GFP(tandem), indicating efficient restoration of surface TCR in the knockout/re-expression system (Figure 1f). In summary, our data shows that the single retroviral vector system allows efficient gene knockout and re-expression in primary murine CD8+ T cells.

### Simultaneous knockout increases integration of ectopically expressed proteins into receptor complexes

We evaluated the advantages of the single vector knockout/re-expression system for analyses of T cell biology. Many immune receptors are multimeric, and competition of ectopically with endogenously expressed subunits can reduce their integration into receptor complexes. This could affect analyses of receptor (i.e. TCR) structure and functions when mutant or tagged subunits are expressed. Thus, we quantified the integration of CD3γ-GFP into cell surface TCR complexes in the absence and presence of endogenous CD3γ. T cell activation by antigen presenting cells (APCs) results in the formation of TCR microclusters at the contact site, known as the immunological synapse ^31,32^. We used ligand-coated surfaces as ‘artificial APCś to analyze CD3γ-GFP recruitment to TCR microclusters.

OT-I Cas9 T cells were transduced with either CD3γ-GFP(mock) or CD3γ-GFP(tandem) to compare CD3γ-GFP integration into TCR in the presence or absence of endogenous CD3γ, respectively. Total surface TCR was detected with fluorescently labeled anti-TCRβ H57 antibody Fab fragments (H57 Fabs). H57 Fabs do not activate T cells and do not interfere with TCR/pMHC interaction^33^. The stained T cells were activated either on glass surfaces coated with antigenic pMHC (H2-K^b^/OVA) (Figure 2a), or OKT3 anti-CD3ε antibody (Supplementary Figure 2). TCR microclusters within the plasma membrane were visualized by Total Internal Reflection Fluorescence Microscopy (TIRFM). The ratio of CD3γ-GFP to H57 Fabs fluorescence was determined to quantify CD3γ-GFP integration into TCR complexes. Based on the substantial loss of endogenous CD3γ in the knockout T cells (Figure 1), the integration of CD3γ-GFP into the TCR in knockout T cells must be close to complete and was therefore normalized to 100%. Integration of CD3γ-GFP into the TCR/CD3 complex was 2- to 3-fold higher in knockout T cells compared to T cells expressing endogenous CD3γ (Figure 2a and 2b, Supplementary Figure 2a and 2b). This increase in integration was not due to differences in CD3γ-GFP or surface TCR expression. CD3γ-GFP expression levels in transduced T cells were approximately 1.5-fold higher when endogenous CD3γ was knock-out (Supplementary Figure 2c). TCR surface expression levels were comparable in all T cells populations (Supplementary Figure 2d). These data demonstrate that studies of signaling complexes, such as the TCR, benefit from the knockout of endogenous genes when expressing mutant or tagged proteins of interest.

**Figure 2.**
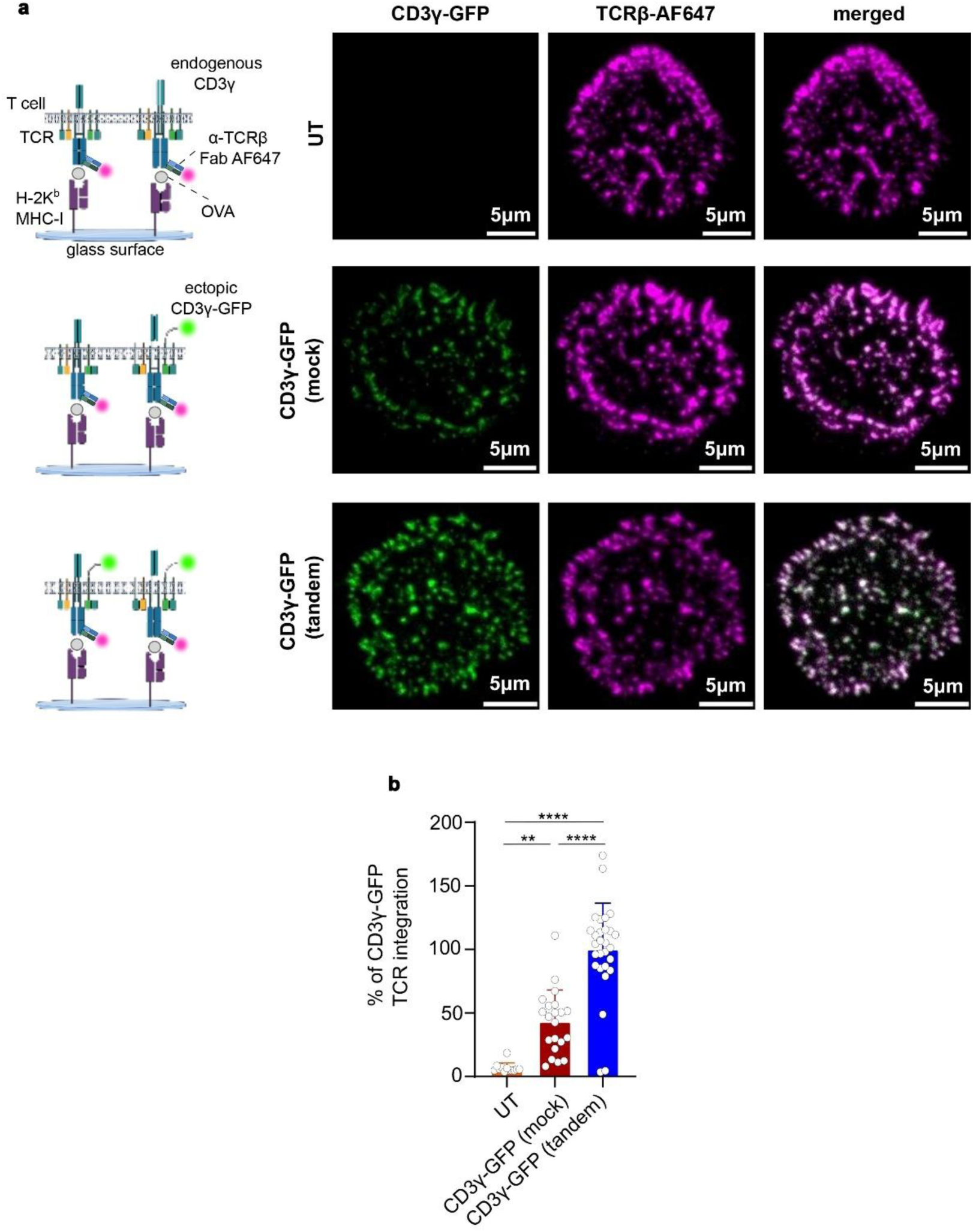
Knockout of endogenous CD3γ increases integration of ectopic CD3γ-GFP into surface TCR complexes. (**a**) (left) Schematic overview of analysis of untransduced (UT), top; CD3γ-GFP(mock), middle; and CD3γ-GFP(tandem)-transduced (bottom) OT-I Cas9 T cells. (right) Representative TIRFM images of transduced OT-I T cells on activating surfaces (H-2K^b^/OVA) showing CD3γ-GFP (left), TCR labelled with AF647-conjugated H57 Fab (TCRβ-AF647; middle) and the merged signals (right column). (**b**) GFP/AF647 pixel intensity ratio normalized to 100% integration for CD3γ-GFP(tandem) sample. Data are presented as mean ± SD, with each dot representing one cell. Statistical analysis was performed using one-way ANOVA with Tukey’s multiple comparisons test (n ≥ 10 cells per sample). Data are presented as mean ± SD. P < 0.05 (*), P < 0.01 (**), P < 0.001 (***), P < 0.0001 (****).

### Knockout and re-expression of cytoplasmic T cell signaling molecules

The tyrosine kinase Zap70 was selected to validate the knockout/re-expression system for functional analyses of different classes of signaling mutants. Zap70 consists of two SH2 (tandem SH2) domains and a kinase domain. The individual SH2 domains are linked by interdomain A, and tandem SH2 is linked to the kinase domain by interdomain B (Figure 3a). The tandem SH2 binds to phosphorylated ITAMs of TCR complex, leading to Zap70 activation and phosphorylation of downstream substrates^24,25^. We developed knockout/re-expression system with a single sgRNA targeting Zap70 and re-expression of HA-tagged WT and mutant Zap70. Two classes of Zap70 mutants were used (Figure 3a): (1) Mutants that compete with endogenous/wild-type Zap70 for binding to the TCR, but do not have kinase activity. Specifically, Zap70 K368A is a full-length protein lacking catalytic activity^34^; and the tandem SH2 domain (tSH2) is a truncated form that lacks interdomain B and the kinase domain^24^. Both mutants bind phosphorylated ITAMs better than wild-type Zap70^24^ and can inhibit T cell activation in Zap70-sufficient T cell lines^34^. Thus, they have been reported as ‘*dominant-negative*’ mutants. (2) A ‘*non-competitive*’ mutant, specifically Zap70 R190/192A^35^, with mutation in the C-terminal SH2 domain that abolishes binding to phosphorylated ITAMs. The phenotype of this mutant is only detectable in Zap70-deficient T cells.

**Figure 3.**
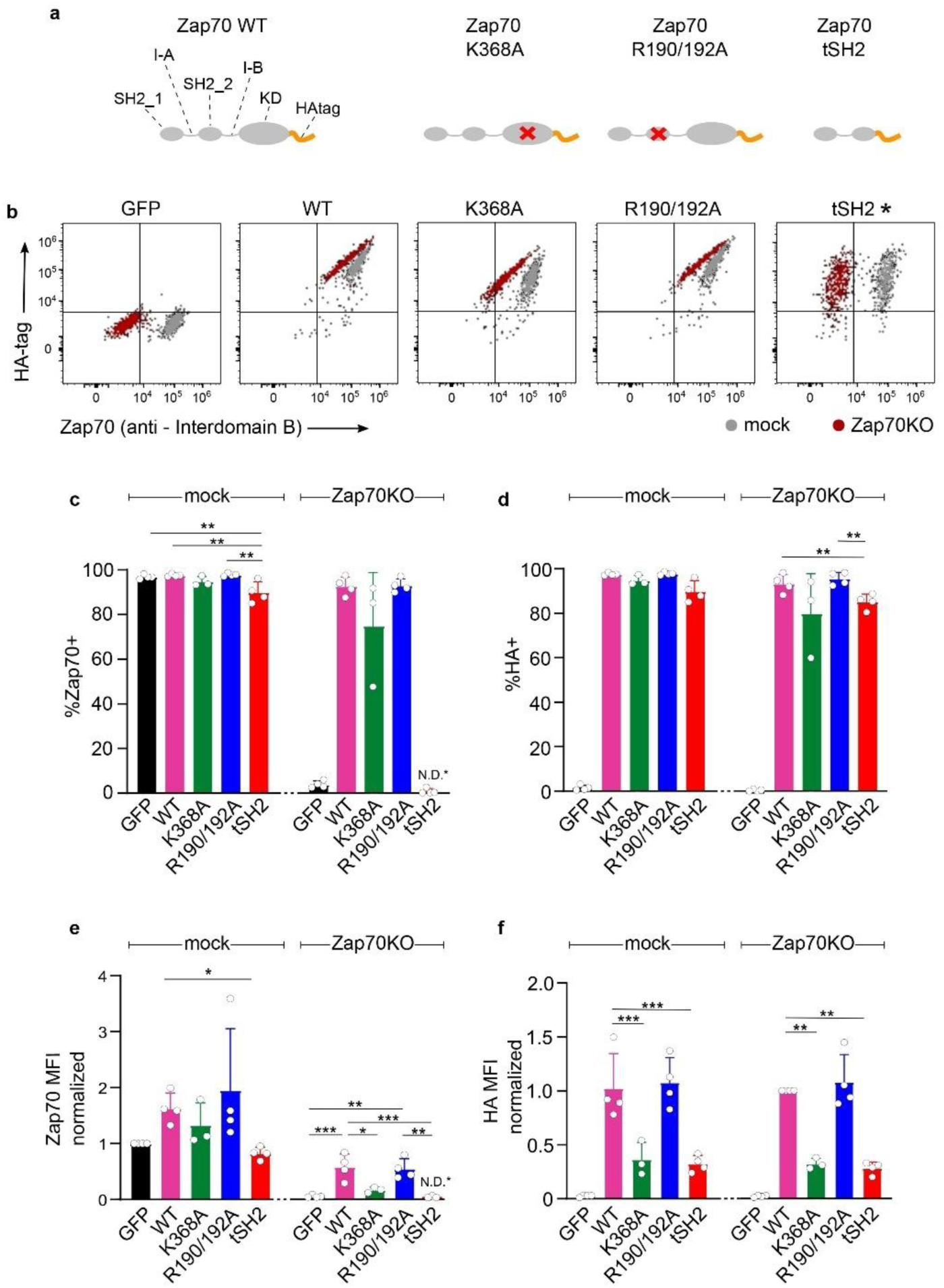
Knockout/re-expression platform to analyze Zap70 mutants in mouse T cells. (**a**) Schematic of Zap70 structure and its mutants. KD = kinase domain, I-A = interdomain A, I-B = interdomain B. (**b-f**) OT-I Cas9 T cells were transduced with constructs encoding mock gRNA (mock) or gRNA targeting Zap70 (Zap70KO) together with the indicated protein (GFP, Zap70 WT or Zap70 mutants), followed by Zap70 and HA intracellular staining on day 7. (**b**) Representative flow cytometry dot plots of total Zap70 (x-axis) and HA-tag (y-axis) of transduced T cells. (tSH2 was not recognized by anti-Zap70 antibody, marked *). (**c+d**) Percentage of Zap70+ and HA+ live cells, respectively. (**e+f**) MFI of Zap70 normalized to GFP(mock) and HA normalized to Zap70(mock), respectively. Data pooled from three to four independent experiments, with one donor mouse per experiment. Data are presented as mean ± SD; in (**c-f**) each dot represents one independent experiment. Statistical analysis was performed using one-way ANOVA with Tukey’s multiple comparisons test. P < 0.05 (*), P < 0.01 (**), P < 0.001 (***).

We expressed wild-type and the three mutant Zap70 in OT-I Cas9 T cells with or without knockout of endogenous Zap70, using sgRNA targeting Zap70 (referred to as Zap70 KO) or mock sgRNA (referred to as mock), respectively (Supplementary Figure 3). Knockout efficiency was tested using a vector that encoded the gRNA and expressed GFP. Total and ectopic HA-tagged Zap70 was quantified using Zap70 and HA intracellular staining on day 7 (Fig. 3b-f). All cells transduced with mock sgRNA remained >95% Zap70 positive (Figure 3b and 3c). Zap70 knockout reduced the Zap70 positive population to less than 5% (Figure 3b and 3c), indicating high knockout efficiency. Re-expression of Zap70 WT, K369A or R190/192A resulted in ∼95%, ∼65% or ∼95% of Zap70 positive T cells, respectively. The re-expression of tSH2 could not be quantified with the anti-Zap70 antibody, as the antibody epitope is not present in the truncated protein. Anti-HA staining allowed detection of all HA-tagged Zap70 variants. Samples transduced with mock gRNA and HA-tagged Zap70 variants contained more than 90% HA-tag positive population (Figure 3d). Knockout and re-expression of Zap70 WT, K369A, R190/192A and tSH2 resulted in ∼95%, ∼70%, 95% and ∼80% HA-tag positive population, respectively, showing good efficiency of knockout and re-expression.

We analyzed Zap70 and HA-tag MFI to asses expression levels of endogenous and ectopic Zap70. Zap70 staining was reduced to background levels in Zap70 knockout samples (Figure 3e). Re-expression of Zap70 WT, K369A and R190/192A resulted in ∼50%, ∼20% and ∼50% of endogenous Zap70 levels, respectively, relative to GFP(mock) control. Expression of these variants without knocking out endogenous Zap70 resulted in staining levels of ∼150%, ∼120% and ∼160%, respectively (Figure 3e). This suggests that expression levels of the endogenous and ectopic Zap70s are additive. For both mock and Zap70 knockout vectors, Zap70 WT and R190/192A were expressed at similar levels (50-60% of endogenous Zap70), while K369A was expressed at about 3-fold lower level (15-20% of endogenous Zap70). HA-tag staining of ectopic Zap70 WT and R190/192A was comparable (Figure 3f), in agreement with data obtained using anti-Zap70 antibody (Figure 3e). Anti-HA staining of Zap70 K369A and tdSH2 were 3-fold lower than WT, indicating that tdSH2 is expressed at 15-20% of endogenous Zap70 levels. The lower expression of Zap70 K369A and tdSH2 is likely due to protein instability and/or increased degradation.

### Phenotypes of non-competitive and dominant negative mutants in primary T cells require gene knockout

Phosphorylated ERK (pERK) was quantified upon TCR ligation to evaluate the knockout/re-expression system for studies of T cell signaling. T cells were stimulated with anti-CD3ε antibody, and pERK levels were monitored over a period of 5 min (Figure 4a). Strong induction of pERK was observed in the mock(GFP) control, peaking at 2 and 3 min with more than 80% pERK positive cells, before declining to 30% at 5 min (Figure 4b). For mock gRNA samples, expression of Zap70 WT or non-competitive R190/192A mutant did not alter the percentage of pERK positive cells or intensity (MFI) of pERK signal (Figure 4b). Unexpectedly, expression of the kinase dead K369A Zap70 mutant did not reduce pERK levels (Figure 4b), suggesting that this mutant does not have the reported dominant negative effects in primary mouse T cells. In contrast, T cells expressing Zap70 tSH2 in presence of endogenous protein showed 2- to 3-fold reduction in percentage of responding T cells, confirming its dominant-negative effect on TCR signaling. These data suggest that lower ectopic expression levels in primary T cells make it less likely to detect dominant-negative effects in the presence of endogenous proteins. The pERK signal was strongly reduced in Zap70 knockout samples, with only 20% pERK positive cells observed (Figure 4b). Moreover, pERK levels were over 2-fold lower compared to mock control. Re-expression of WT Zap70 fully restored pERK signal. However, none of the mutants tested rescued Erk phosphorylation. This is in contrast to unimpaired pERK responses when R190/192A or K369A were expressed in presence of endogenous Zap70, further supporting that their phenotypes are suppressed by endogenous Zap70. This shows the necessity for an effective knockout and re-expression for analysis of both non-competitive and dominant negative mutants in primary T cells.

**Figure 4.**
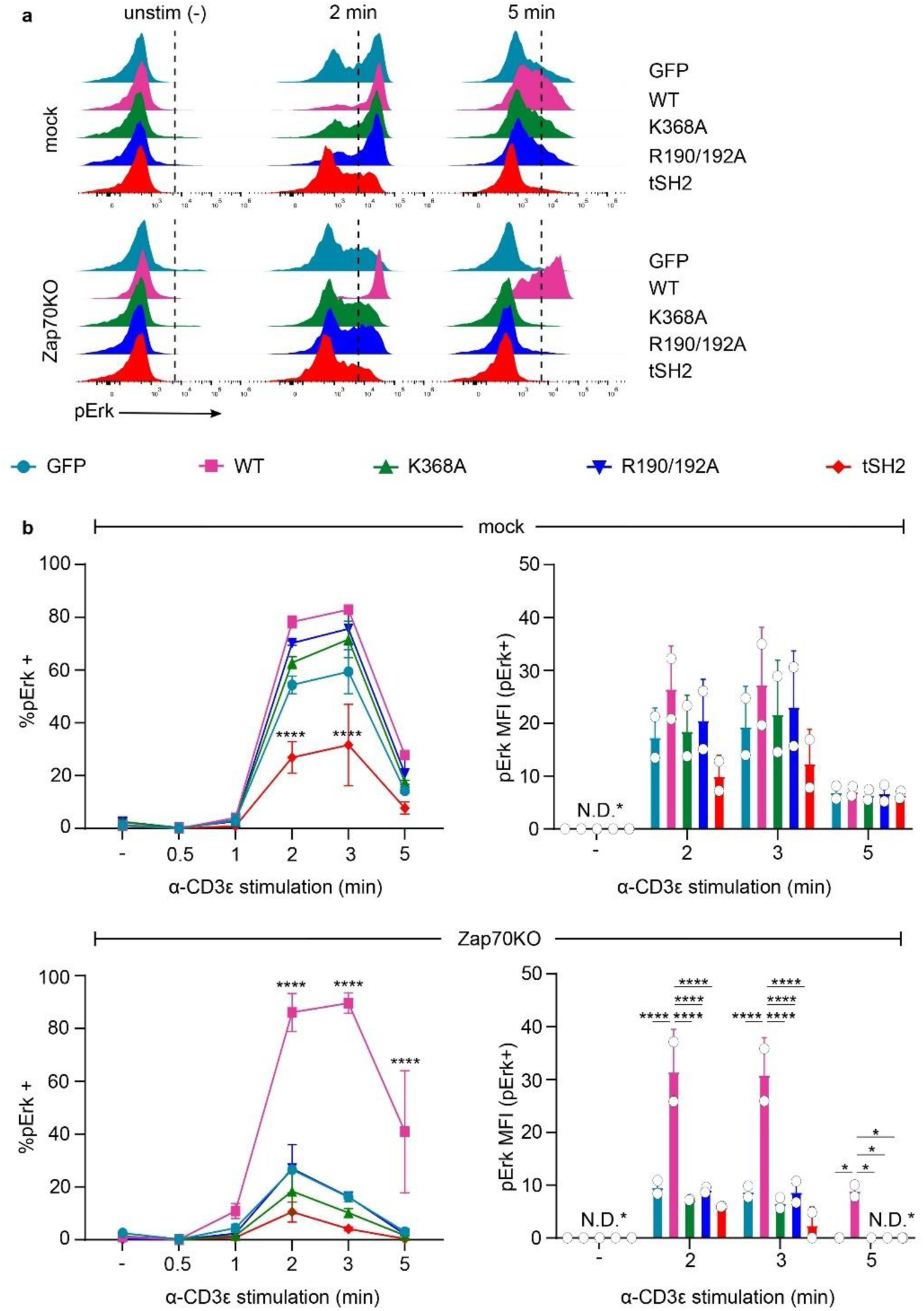
Signaling phenotype of Zap70 mutants requires knockout of endogenous Zap70. OT-I Cas9 T cells were transduced with construct encoding mock gRNA (mock) or gRNA targeting Zap70 (Zap70KO) together with the indicated proteins (GFP, Zap70 WT or Zap70 mutants). On day 7, T cells were stimulated with anti-CD3ε antibody (clone 2C11) for indicated time points. ERK phosphorylation (pERK) was analyzed by flow cytometry. (**a**) Representative flow cytometry histograms showing pERK+ gate. (**b**) Percentage of pERK+ T cells (left) and MFI of pERK in pERK+ T cells (right) transduced with construct encoding mock (top) or Zap70 gRNA (bottom) with ectopically expressed GFP, wild-type or mutant Zap70s. Data pooled from two independent experiments, with one donor mouse per experiment. Data are presented as mean ± SD. Statistical analysis was performed using two-way ANOVA followed by Tukey’s multiple comparisons test. P < 0.05 (*), P < 0.01 (**), P<0.001 (***), P < 0.0001 (****).

We then compared the effects of Zap70 mutants on T cell functions. T cells transduced with mock or Zap70 KO constructs, co-expressing either HA-tagged Zap70 variants (WT, K368A, R190/192A, or tSH2) or GFP, were activated using EL4 target cells pulsed with different concentrations of OVA peptide. Cytokine production was measured using intracellular staining to detect IFNγ and TNFα. Cytotoxicity was quantified using staining to detect CD107a on T cell surface as a marker for cytotoxic granule release, and by directly measuring target cell death. We observed efficient cytokine production, degranulation and target cell killing in control mock(GFP) T cells (Figure 5a-f, Supplementary Figure 4). Expression of any Zap70 mutant with mock gRNA did not affect these responses (Figure 5a, 5c and 5e), indicating that endogenous Zap70 compensates for effects of the mutants in T cell effector function. The unimpaired effector functions of mock cells expressing tdSH2 are in contrast to their reduced pERK responses (Figure 4), suggesting stronger compensatory mechanisms for induction of T cell functions as compared to signaling. Zap70 knockout abolished cytokine production, degranulation and target cell killing, which were restored by expression of WT Zap70, but not any of the mutants tested (Figure 5b, 5d and 5f, Supplementary Figure 4). To account for different expression levels of Zap70 mutants, we analyzed cytokine production in Zap70 knockout populations with matching ectopic Zap70 levels, based on anti-HA staining (Supplementary Figure 5a). Low levels of ectopic WT Zap70 restored cytokine production, but even high levels of any of the mutants did not (Supplementary Figure 5b and 5c). This indicates that the observed reduction in T cell effector functions reflect the phenotype of Zap70 mutants independently of their expression levels. Overall, our results show that knock-out of endogenous Zap70 is required to reveal effector phenotypes of both non-competitive and dominant-negative mutants in primary mouse T cells.

**Figure 5.**
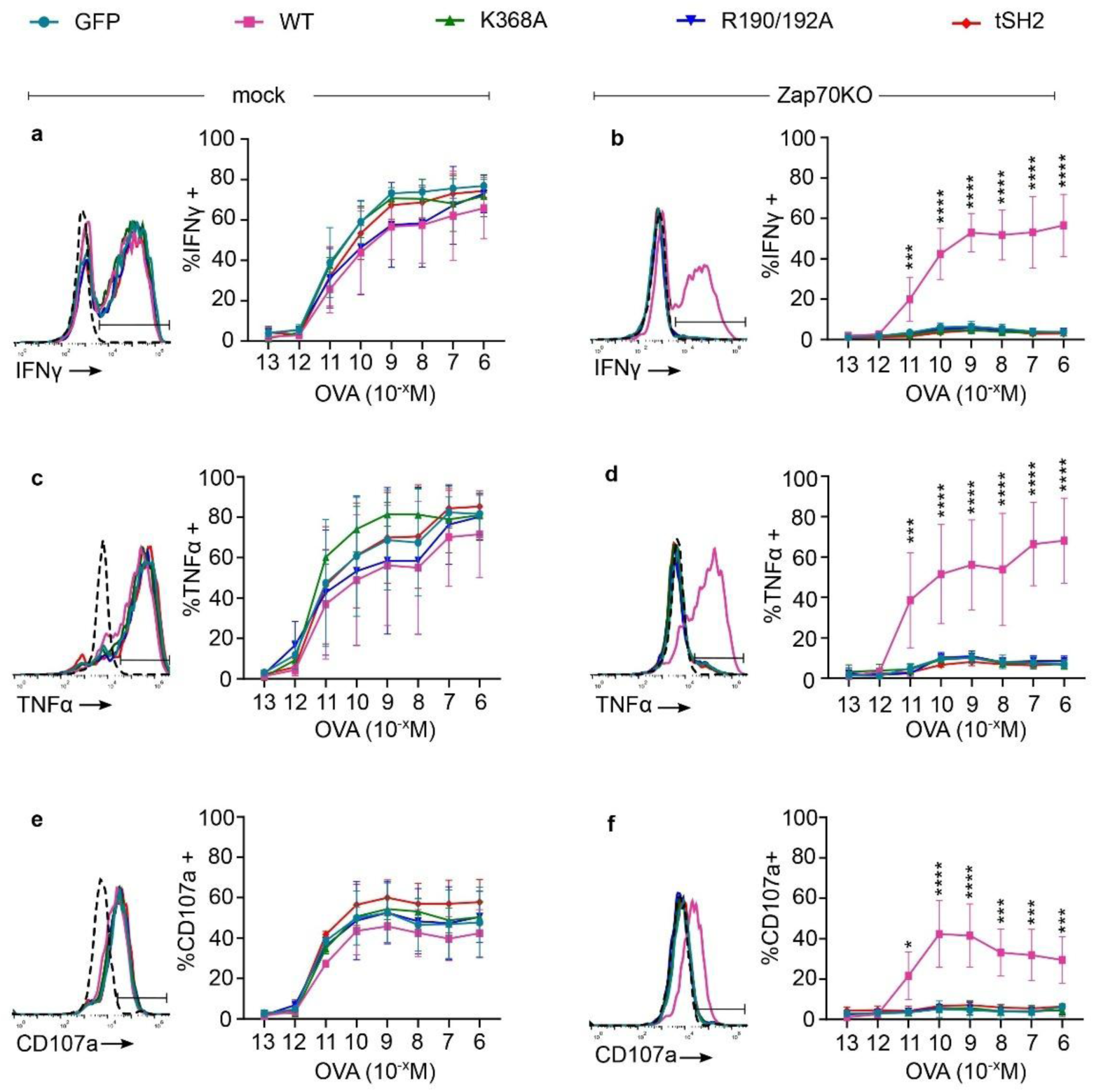
Effector phenotype of Zap70 mutants requires knockout of endogenous Zap70. OT-I Cas9 T cells were transduced with construct encoding mock gRNA (mock) or gRNA targeting Zap70 (Zap70KO) together with the indicated proteins (GFP, Zap70 WT or Zap70 mutants). EL4 target cells were loaded with different concentrations of the high affinity N4 peptide and used to stimulate transduced T cells. Intracellular staining was used to quantify (**a+b**) IFNγ and (**c+d**) TNFα production, (**e+f**) surface staining for CD107a was used as a measure of degranulation. Percentage of IFNγ, TNFα or CD107a positive cells are shown for mock (**a,c,e**) or Zap70KO (**b,d,f**) transduced T cells. Representative flow cytometry histograms (left) are show for 10^−9^M OVA peptide, and include unstimulated control (dotted black line). Data pooled from three independent experiments, with one donor mouse per experiment. Data are presented as mean ± SD. Statistical analysis was performed using two-way ANOVA followed by Tukey’s multiple comparisons test. P < 0.05 (*), P < 0.01 (**), P < 0.001 (***), P < 0.0001 (****).

### Knockout/re-expression platform to analyze mutants *in vivo*

Finally, we evaluated the knockout/re-expression system’s benefits for analyzing anti-tumor T cell responses *in vitro* and *in vivo*. We analyzed the non-competitive R190/192A mutant in the absence and presence of endogenous Zap70, as it requires the most efficient depletion of endogenous Zap70. Murine MC38 adenocarcinoma was used as a model tumor cell line. MC38 cells were modified to express cytoplasmic variant of ovalbumin, ensuring physiological processing and presentation of antigenic OVA peptide (MC38-OVA cell line). As a control, the OVA peptide sequence within ovalbumin was replaced with that of null (SIAAFASL^36^) peptide (MC38-null cell line). OT-I Cas9 T cells were transduced with either GFP(mock or Zap70KO) or Zap70 R190/192A(mock or Zap70KO), and tested *in vitro* and *in vivo* using the MC38 tumor model.

Cytokine production and CD25 upregulation were analyzed to quantify anti-tumor T cell responses *in vitro*. Control MC38-null cells did not induce cytokine secretion or CD25 upregulation (Figure 6a-f). GFP(mock) T cells expressing endogenous Zap70 secreted cytokines and upregulated CD25 in response to MC38-OVA tumor cells. This response was not altered by ectopic expression of Zap70 R190/192A (Figure 6a-f). Zap70 knockout abolished responses to antigenic tumor cells, which was not restored by ectopic expression of R190/192A mutant. These findings agree with our data using anti-CD3 stimulation (Figure 4) and peptide-pulsed target cells (Figure 5), showing that the phenotype of R190/192A ZAP70 becomes apparent only in the absence of endogenous Zap70.

**Figure 6.**
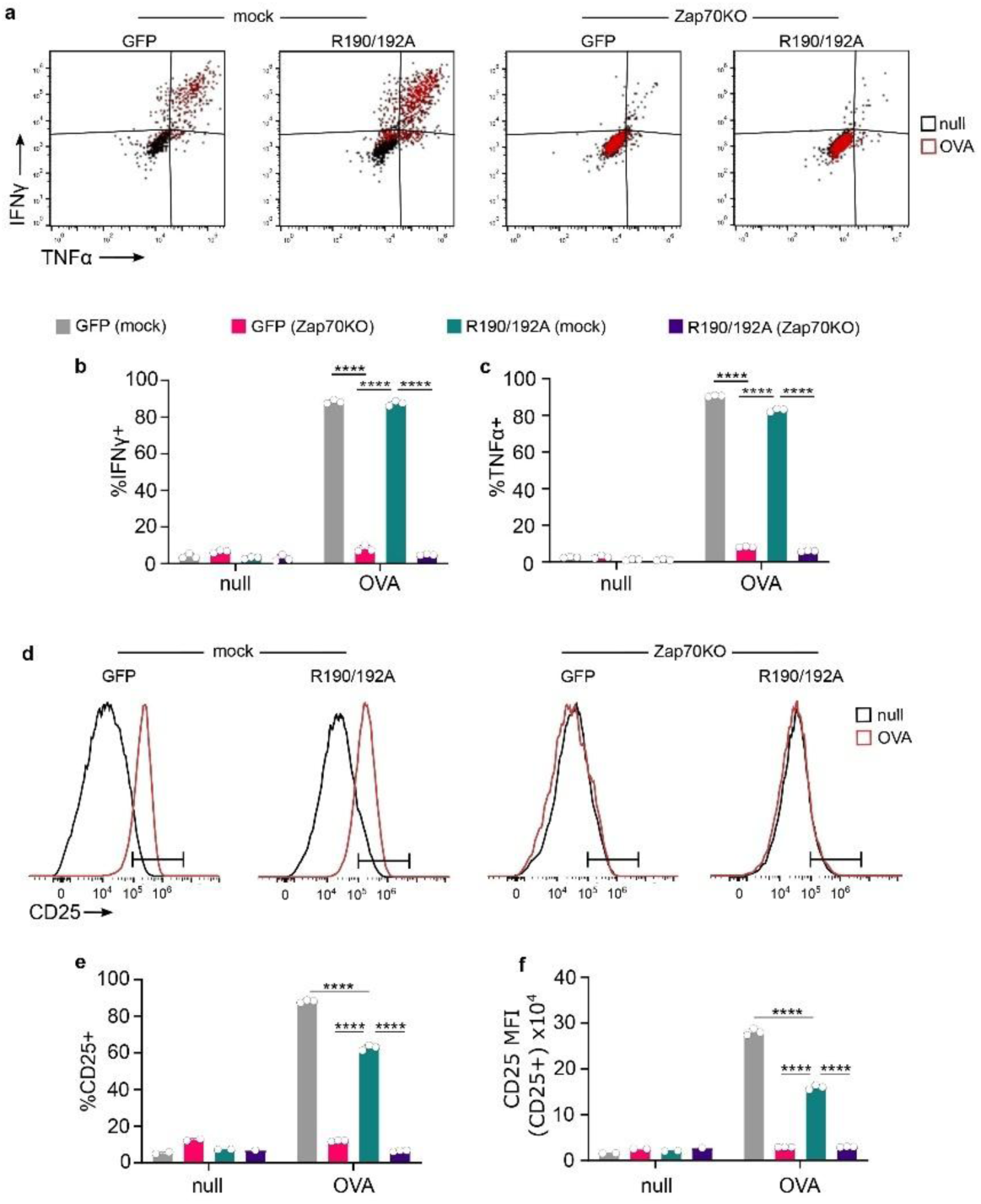
Single vector knockout/re-expression system supports *in vitro* analysis of anti-tumor T cell responses. OT-I Cas9 T cells were transduced with construct encoding mock gRNA (mock) or gRNA targeting Zap70 (Zap70KO) together with either GFP or Zap70 R190/192A. Transduced T cells were co-cultured with MC38 cells expressing null or OVA peptide. (**a-d**) Cytokine production was measured using flow cytometry after 5h co-culture. (**a**) Representative flow cytometry plot of TNFα (x-axis) and IFNγ (y-axis). (**b+c**) Percentages of IFNγ+ and TNFα+ CD8+ T cells, respectively. (**d**) CD25 surface expression of transduced T cells was measured using flow cytometry after 24h co-culture. Representative flow cytometry histograms showing CD25+ gate. (**e**) Percentages of CD25+ CD8+ T cells. (**f**) CD25 MFI of CD25+ T cell population. Data from one experiment, with 3 technical replicates per condition. Data are presented as mean ± SD; in (c-f) each dot represents one replicate. Statistical analysis was performed using two-way ANOVA with Sidak’s multiple comparison. P < 0.0001 (****).

CD8 T cells can directly kill tumor cells, and this cytotoxic function is critical for control of tumor growth. We measured target cell death (Figure 7a) and CD8 T cell degranulation (Figure 7b, 7c and 7d) to evaluate the effect of Zap70 R190/192A expressed in presence or absence of endogenous Zap70 on anti-tumor cytotoxicity. GFP(mock), but not GFP(Zap70 KO) T cells degranulated and killed antigenic tumor cells (Figure 7a-d). Ectopic expression of R190/192A mutant had no effect on cytotoxicity of mock-transduced cells, but failed to restore cytotoxicity of Zap70 knockout cells.

**Figure 7.**
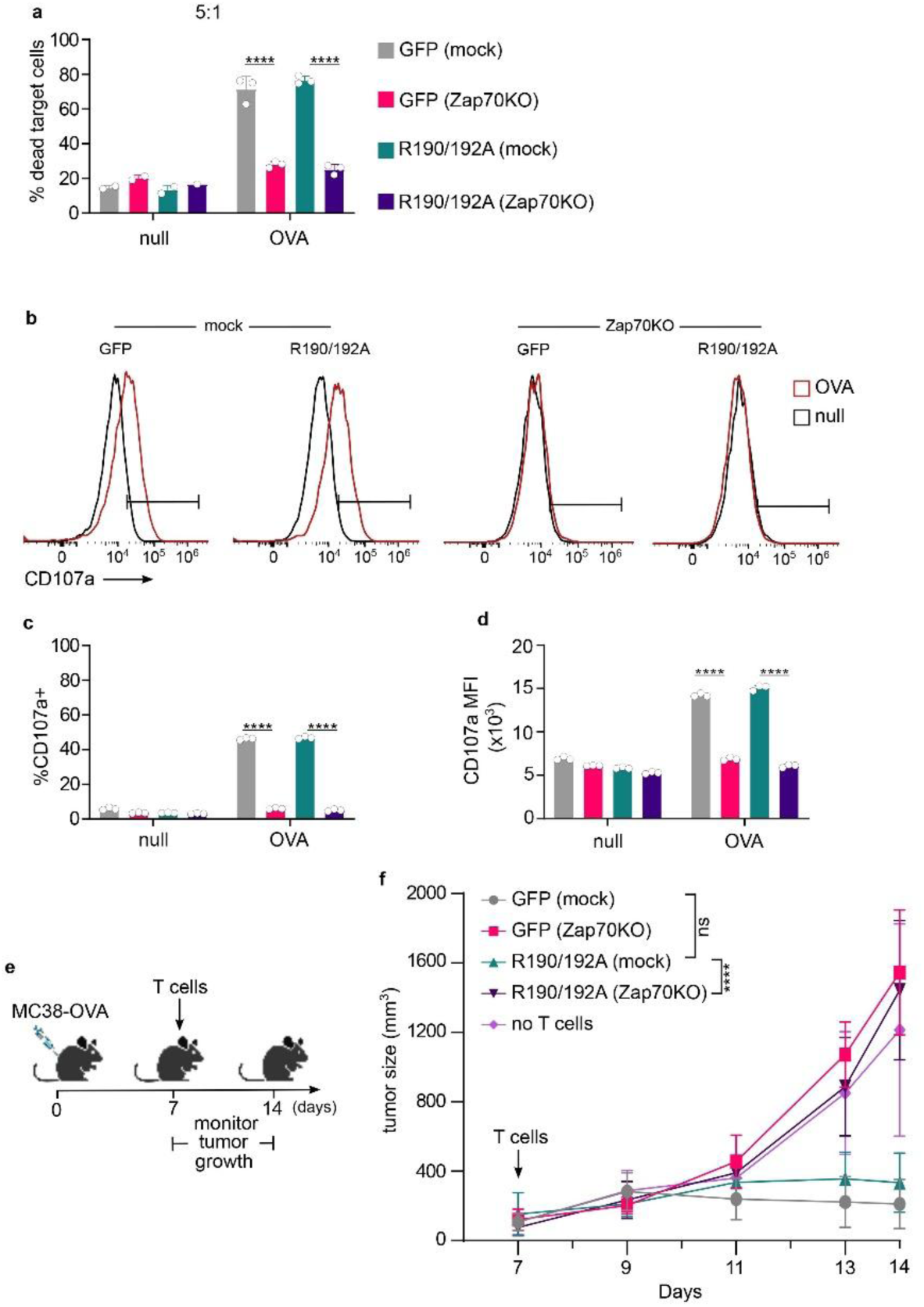
Single vector knockout/re-expression system supports in vivo analysis of anti-tumor T cell responses. OT-I Cas9 T cells were transduced with construct encoding mock gRNA (mock) or gRNA targeting Zap70 (Zap70KO) together with GFP or Zap70 R190/192A. Cytotoxic responses against MC38 cells expressing null or OVA peptide were measured. (**a**) T cells were incubated with MC38 cells for 24h at indicated effector to target (E:T) ratios. Percentage of dead MC38 cells was measured using flow cytometry. (**b**) T cells were incubated with MC38 cells for 5h, and CD107a staining was measured. Representative flow cytometry histogram showing CD107a+ gate used. (**c**) Percentages of CD107a+ CD8+ T cells. (**d**) CD25 MFI of the CD8+ T cell population. (**e**) Timeline of the analysis of anti-tumor T cell responses. MC38 cells expressing null or N4 were subcutaneously injected into Rag1KO recipients, followed by adoptive transfer of T cells transduced with construct encoding mock gRNA (mock) or gRNA targeting Zap70 (Zap70KO) together with GFP or Zap70 R190/192A. (**f**) Tumor volumes of mice injected with differently transduced T cells. The control group did not receive T cells (no T cells). (**a-d**) Data from one experiment, with 2-3 replicates per condition. Data are presented as mean ± SD; in (a,c,d) each dot represents one replicate. (**e+f**) Data from one experiment, using 5 donor mice per group. Statistical analysis was performed using two-way ANOVA followed by Sidak’s multiple-comparisons test (a-d) or Tukey’s multiple-comparisons test (f). ns = not significant, P < 0.0001 (****).

Anti-tumor T cell responses *in vivo* depend on several factors, such as cell trafficking, survival and differentiation, which are not fully recapitulated using *in vitro* models. However, *in vivo* mutant analysis often requires pure gene edited population, as presence of small fraction of wild-type or non-edited T cells can influence tumor control. Obtaining sufficiently pure gene-edited populations is challenging, especially for intracellular signaling proteins, when the desired cell populations cannot be sorted based on surface proteins expression. However, the high knockout and re-expression efficiencies obtained in our system offer possibility of overcoming this limitation. We therefore tested if our single vector knockout/re-expression platform can be used for analysis of anti-tumor T cells responses *in vivo*. MC38-OVA tumor cells were injected subcutaneously into immunodeficient Rag1KO recipients, and transduced OT-I Cas9 T cells were adoptively transferred 7 days later (Figure 7e). We observed tumor progression in control mice that did not receive T cells, but GFP(mock) T cells controlled tumor growth (Figure 7d). Zap70 knockout T cells failed to control tumors, in agreement with our results from *in vitro* experiments (Figure 7a-d). Zap70 knockout T cells re-expressing Zap70 R190/192A did not control tumor growth, in contrast to R190/192A(mock) cells (Figure 7d). This confirms our previous finding that the phenotype Zap70 R190/192A is apparent only in the absence of endogenous Zap70, and underscores the benefits of this knockout/re-expression systems for mutant analysis during anti-tumor responses.

### Two-vector system for knock-out and re-expression in Cas9-deficient primary T cells

Many mouse models are not available as Cas9 transgenics, and could not be analyzed with our single vector system. Therefore, we tested if our knockout/re-expression vector can be combined with a second vector encoding Cas9 for analyses of additional mouse models. Although Cas9 (∼4.1 kb) is within packaging limits for γ-retroviruses (7-8kb), its relatively large size can reduce packaging efficiency. Indeed, use of a single retroviral vector encoding Cas9 and sgRNA in mouse T cells was reported in result in low transduction (30-60%) and knock-out efficiencies (15-50% of the transduced cells)^1^. We optimized our alternative two-vector system for high transduction and editing efficiency. We designed a retroviral vector encoding Cas9 and blasticidin resistance gene joined via T2A self-cleaving peptide. T2A and blasticidin resistance were chosen due to their relatively small size. The knockout/re-expression vectors contained puromycin resistance gene to allow antibiotic selection of the doubly transduced T cell. Specifically, blastidicine resistance gene in the Zap70 knockout/re-expression vectors described before (Supplementary Figure 3) was replaced with that of puromycin. The packaging cells were separately transfected with the knockout/re-expression vector or Cas9 vector. The resulting supernatants were pooled at 1:3 knockout/re-expression:Cas9 ratio to enhance Cas9 transduction efficiency.

Transduction of OT-I T cells with Cas9 and GFP(tandem) targeting CD3γ resulted in efficient CD3γ knockout, with more than 90% of the transduced cells showing loss of surface TCR (Figure 8a). CD3γ-GFP(tandem) vector fully restored TCR surface expression (Figure 8a), indicating that our system can be used to knockout and re-express proteins in T cells from any mouse strain. We then tested if this approach can be used to modify cytoplasmic proteins in T cells, by targeting the kinase Zap70.

**Figure 8.**
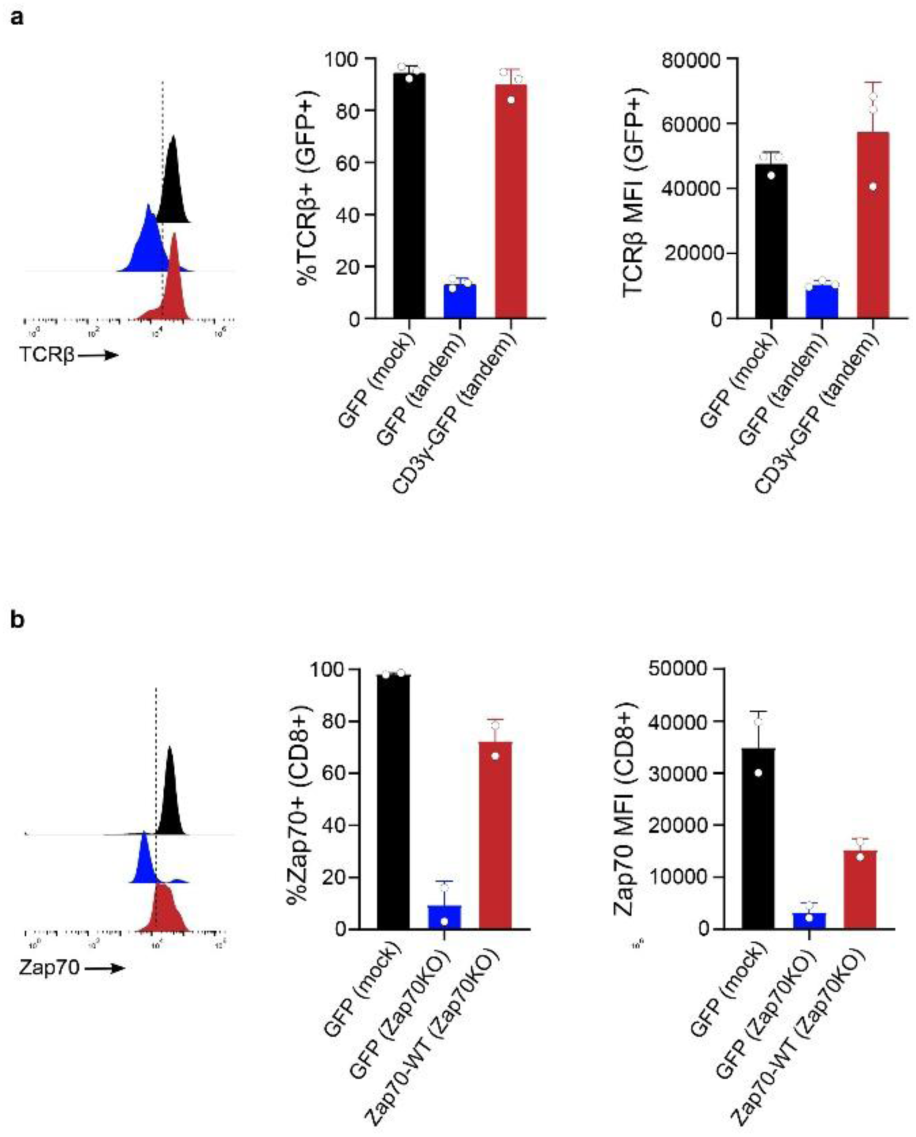
Two-vector system for knock-out and re-expression in Cas9-deficient primary T cells. (**a**) OT-I T cells were transduced with a vector encoding Cas9 and a vector encoding mock gRNA (mock) together with GFP, or gRNA targeting CD3γ (tandem) together with GFP or CD3γ-GFP. Representative flow cytometry histograms showing surface TCRβ staining in GFP+ cells. Graphs show percentage of TCRβ+ T cells within the GFP+ population (left) and TCRβ MFI of GFP+ cells (right). (**b**) OT-I T cells were transduced with a vector encoding Cas9 and a vector encoding mock gRNA (mock) together with GFP, or gRNA targeting Zap70 (Zap70KO) together with GFP or HA-tagged Zap70-WT. Representative flow cytometry histograms showing Zap70 staining in CD8+ T cells. Graphs show percentage of Zap70+ T cells within the CD8+ population (left) and Zap70 MFI of CD8+ cells (right). Data from 2-3 independent experiments with one donor mouse per experiment are presented as mean ± SD.

Transduction of OT-I T cells with Cas9 and GFP(Zap70 KO) constructs resulted in efficient Zap70 knockout, with more than 90% of the transduced cells showing loss of Zap70 (Figure 8b). Use of Zap70-WT(Zap70 KO) construct restored Zap70 expression in ∼70% of the transduced cells. Although we observed a lower recovery for Zap70 than CD3γ, both expression levels are sufficient for mutant analysis, especially when combined with staining to detect the ectopically expressed proteins (Supplementary Figure 5). T cell numbers increased approximately 2-fold over four days after transduction (day 7). Thus, knockout and re-expression in T cells from any mouse strain is proficient using an optimized Cas9 vector with our knockout/re-expression system.

## Discussion

We developed a retroviral system that enables highly efficient CRISPR/Cas9-mediated knockout and simultaneous re-expression of wild-type or mutant proteins in T cells from both Cas9 transgenic or non-transgenic mice. This approach leverages the efficiencies of retroviral transduction and CRISPR/Cas9 gene editing for analysis of mutant phenotypes. We validated the method by targeting CD3γ and Zap70, as examples of transmembrane and cytoplasmic T cell signaling proteins, respectively. We observed robust knockout for both, which was rescued by protein re-expression. We have shown that knockout of endogenous protein (i.e. CD3γ) enhances integration of its ectopically expressed tagged version into signaling complexes. We validated our system for functional analysis of mutant phenotypes using well characterized of Zap70 mutants. Zap70 knockout abolished T cell activation, which was rescued by expression of WT Zap70, but not mutants deficient in phospho-ligand binding (non-competitive mutants) or kinase activity (dominant negative mutants). Most importantly, we have shown that our system can be used to generate gene-edited primary T cells for analysis of anti-tumor responses *in vivo*. Our approach expands the toolbox for gene editing in primary murine T cells, by using retroviral transduction for comprehensive analysis of mutant phenotypes.

Our platform provides a practical alternative to knockin T cells for mutant analysis. It has the advantage of high editing efficiency and ease of implementation for any laboratory that uses retroviruses to transduce mouse T cells. Mouse T cell knockin can be achieved by electroporation, but this reduces cell viability and has low knockin efficiency^10,13^. Recent developments in AAV-based gene editing have significantly improved knockin efficiencies in primary mouse T cells^16^. AAV-based delivery of gRNA and repair templates into Cas9-expressing T cells generates knockins in up to 50% cells, or up to 75% when NHEJ inhibitor is used. However, unlike retroviral transduction, this method is not implemented in most laboratories working on murine lymphocytes. Importantly, our approach can be used in T cells that do not express Cas9. The relatively large size of Cas9 has been considered a limiting factor for retroviral transduction^1^. Here, we show that Cas9 vector can be delivered together with knockout/re-expression vector for efficient knockout and re-expression. This extends the utility of our system to any mouse strain without the need for breeding with Cas9 transgenic mice.

We also tested if gRNA multiplexing can be used enhance knockout efficiency in mouse T cells. The multiplexing is based on use of tRNA to connect gRNA sequences^26,27^. Two gRNAs targeting CD3γ were used as a model system to compare knockout efficiency of single and multiplexed (tandem) gRNAs. Each individual gRNA downregulated TCR in approximately 90% of transduced cells, whereas the tandem gRNA downregulated TCR in almost all cells. This result indicates that gRNA multiplexing can be used to improve knockout efficiencies in primary mouse T cells. Similar multiplexing approach was previously used in mouse T cells^37,38^. However, the knockout efficiencies were not reported, or compared between single and tandem gRNA constructs. gRNA multiplexing to improve knockout efficiency can be of special interest when investigating functions of intracellular proteins, when pure knockout populations cannot be obtained by cell sorting based on surface staining.

The knockout/re-expression platform established here provides a powerful approach for mutant analysis in primary murine T cells *in vitro* and *in vivo*. Nevertheless, there are certain limitations of the system. Our approach is based on retroviral transduction, making it easy to implement in most laboratories working on mouse T cells. However, this limits its use to analysis of activated, but not naïve T cells. The system provides a very efficient alternative to knockin for analysis of mutant phenotypes. However, expression of the gene of interest is not controlled by endogenous regulatory mechanisms, which could be a confounding factor for certain phenotypes. This possible limitation can be offset by promoter choice and/or analysis of sub-populations with defined expression levels. Our system can be used for analysis of anti-tumor T cell functions *in vivo*. However, Cas9 protein, proteins encoded by antibiotic resistance genes or protein-tags can be immunogenic^39^. This is a common problem when gene edited T cell are used for adoptive transfers, and it can be offset by use of immunodeficient or immunologically tolerant recipients^40^ or by reducing transgene immunogenicity^41^.

Protein complexes play a critical role in T cell signaling. Their assembly, spatial arrangement and post-translational modifications are investigated using mutated or tagged versions of ectopically expressed proteins. However, the presence of endogenous protein can decrease incorporation of these mutated or tagged proteins into signaling complexes. This can increase their detection thresholds (e.g. for microscopy) and/or mask their effects on cellular functions (e.g. signal transduction). Similar problems exist for the analyses of many cellular structures and protein assemblies/networks. Knockin mice expressing T cell signaling proteins can be generated^42–44^, but this approach is rare due to the cost and time required. Here, we analyzed the incorporation of CD3γ-GFP into the TCR complex to demonstrate that our knockout/re-expression system is ideally suited for the structural and functional analyses of ectopically expressed proteins.

We validated of our knockout/re-expression platform for analysis of mutant phenotypes using previously characterized Zap70 mutants^24,34^. We compared the phenotypes of dominant negative (i.e. K368A and tSH2) or non-competitive (i.e. R190/192A) Zap70 mutants. All mutants showed full or partial T cell activation in the presence of endogenous Zap70 in an array of cellular assays. However, their complete negative phenotype was easily observed when endogenous Zap70 was removed using our knockout/re-expression system. This is because knockout of endogenous protein eliminates competition between wild-type/endogenous and mutant proteins. This suggests that many reported dominant negative effects only result from high levels of overexpression in T cell lines. Indeed, the dominant negative phenotype of two mutants used here (i.e. K368A and tSH2) were shown to be dose-dependent in Jurkat T cells^34^. Our data demonstrates that our retroviral knockout/re-expression system can reveal phenotypes of both non-competitive and dominant negative mutants, while avoiding artifacts associated with the use of cell lines.

Knockout and mutant analyses are widely used in pre-clinical models of T cell-based immunotherapies. These rely on CRISPR/Cas9-mediated knockouts^45,46^, ectopic expression of mutants ^47^ or use of knockin mice^48^. We tested our knockout/re-expression system for analysis of anti-tumor responses *in vitro* and *in vivo*, focusing on the non-competitive Zap70 R190/192A mutant. Zap70-suffcient T cells showed strong anti-tumor effector functions *in vitro*, and controlled tumor growth *in vivo*. Anti-tumor responses of Zap70-suffcient T cells were not affected by expression of the R190/192A mutant. Knockout of Zap70 abolished anti-tumor T cell responses, which were not rescued by expression of the R190/192A mutant. These findings highlight the utility of our knockout/re-expression system for analysis of mutant phenotypes, especially when non-competitive mutants are masked by the presence of endogenous protein. The high knockout and re-expression efficiency achieved in our system allows adoptive transfers without additional sorting for edited populations, which is especially useful for analyses of cytoplasmic proteins. Moreover, the high efficiency is an advantage when analyzing gene edited T cells with impaired proliferation or differentiation, which can be outcompeted by WT T cells^49^.

In conclusion, our knockout/re-expression system provides a powerful tool for gene editing in primary murine T cells. Currently, most of mutant analyses in T cells rely on ectopic expression in knockout cell lines or generation of knockin mice. Our system bridges the two approaches, allowing straightforward, fast, efficient and low-cost mutant analysis in primary mouse T cells.

## Acknowledgements

This study was supported by the Deutsche Forschungsgemeinschaft (DFG, German Research Foundation) through Art. 91B GG funding (INST 39/1323-1 FUGG, INST 39/1324-1 FUGG and INST 39/1325-1 FUGG) and under Germany’s Excellence Strategy (CIBSS - EXC-2189 - Project ID 390939984); A Struktur- and Innovationsfonds für die Forschung (SIBW-Projekt: 7100025401); and The National Institute of Health (NIH 5R01GM118879). We would like to thank the NIH tetramer core facility (Emory University) for providing H-2K^b^OVA monomers. J.B. was supported by Innovation Fund Research 2026/2 – Funding line START from Freiburg University. V.M. was supported by the EURIdoc programme, funded by the European Union’s Horizon 2020 research and innovation programme under the Marie Skłodowska-Curie grant agreement number 101034170.

---

**Supplementary Figure 1.**
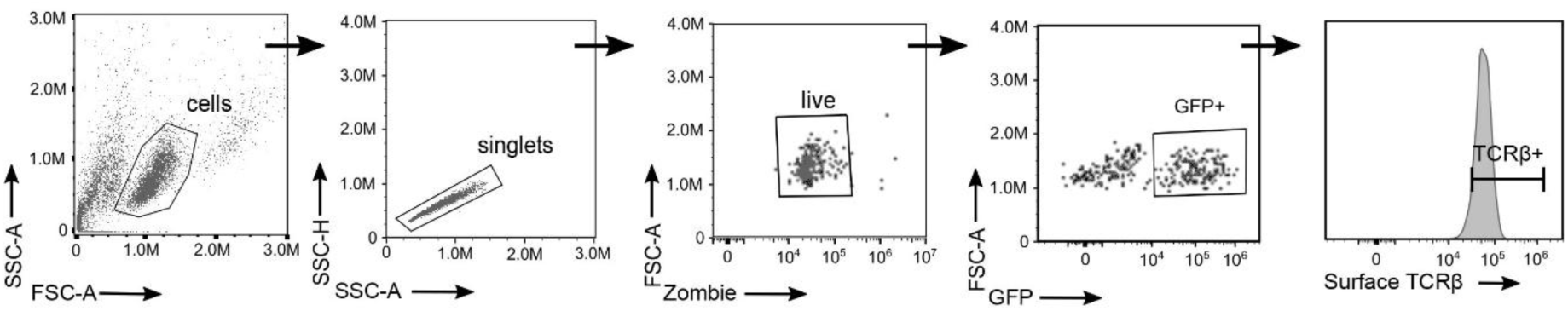
Single vector system to knockout and re-express CD3γ in mouse T cells. Gating strategy used for analysis presented in Figure 1.

**Supplementary Figure 2.**
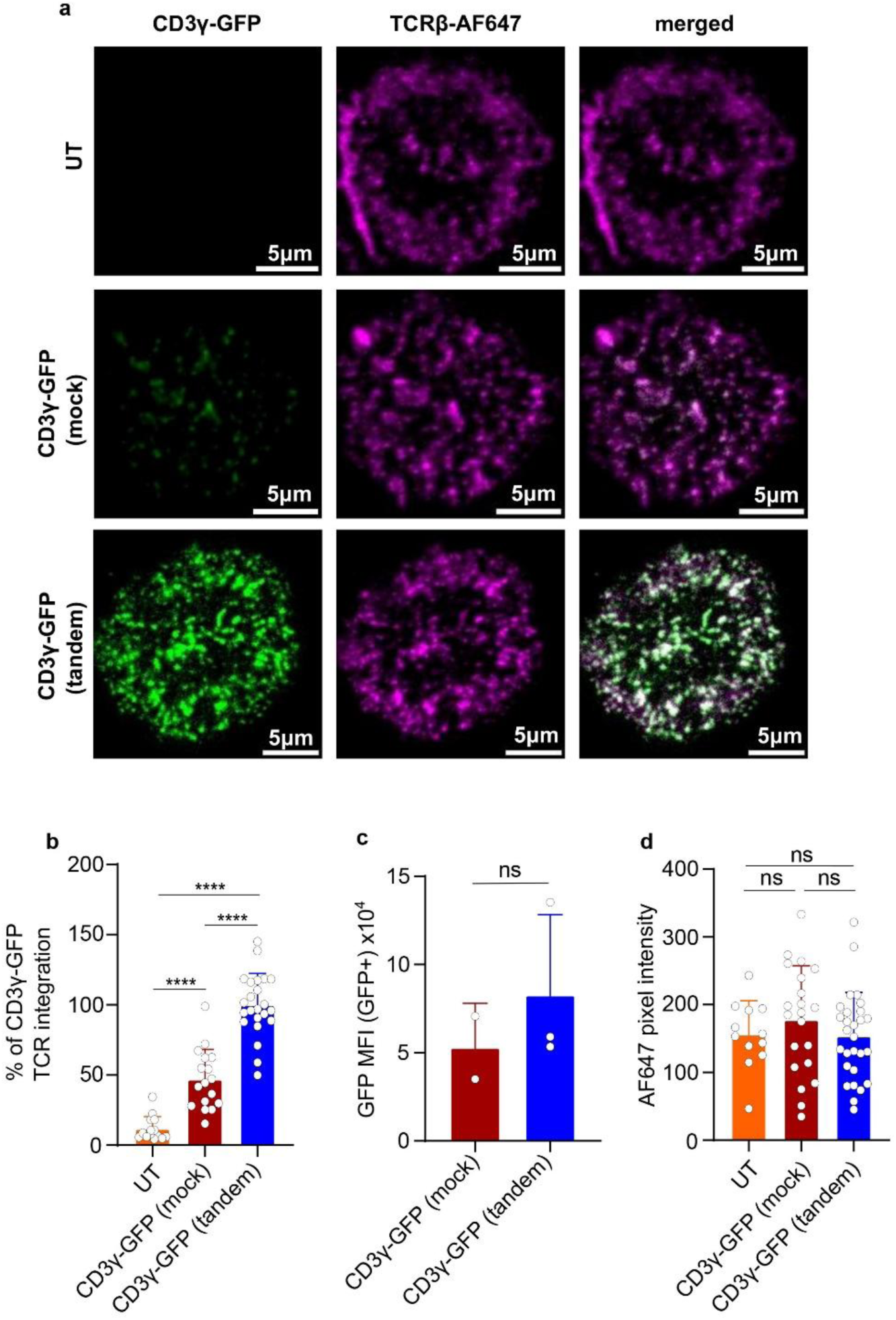
Endogenous CD3γ reduces integration of ectopic CD3γ-GFP into surface TCR complexes. (a) Untransduced (UT), CD3γ-GFP(mock) and CD3γ-GFP(tandem)-transduced OT-I Cas9 T cells were stimulated on coverslips coated with OKT3 anti-CD3ε antibody. Representative TIRFM images of T cells for CD3γ-GFP (left), TCR labelled with AF647-conjugated H57 Fab (TCRβ-AF647, middle) and the merged signal (right). (b) GFP/AF647 pixel intensity ratio normalized to 100% integration for CD3γ-GFP(tandem) samples on OKT3 surfaces. (c) Flow cytometry analysis of CD3γ-GFP expression levels in CD3γ-GFP(mock) and CD3γ-GFP(tandem) transduced T cells for Figure 2 and Supplementary Figure 2. (d) AF647 (H57 Fab anti-TCR) pixel intensity used to calculate AF647:GFP ratio in Figure 2b and Supplementary Figure 2. Data are presented as mean ± SD, with each dot representing one cell. Statistical analysis was performed using one-way ANOVA with Tukey’s multiple comparisons test (n ≥ 10 cells per sample). Data are presented as mean ± SD. ns = not significant, P < 0.0001 (****).

**Supplementary Figure 3.**
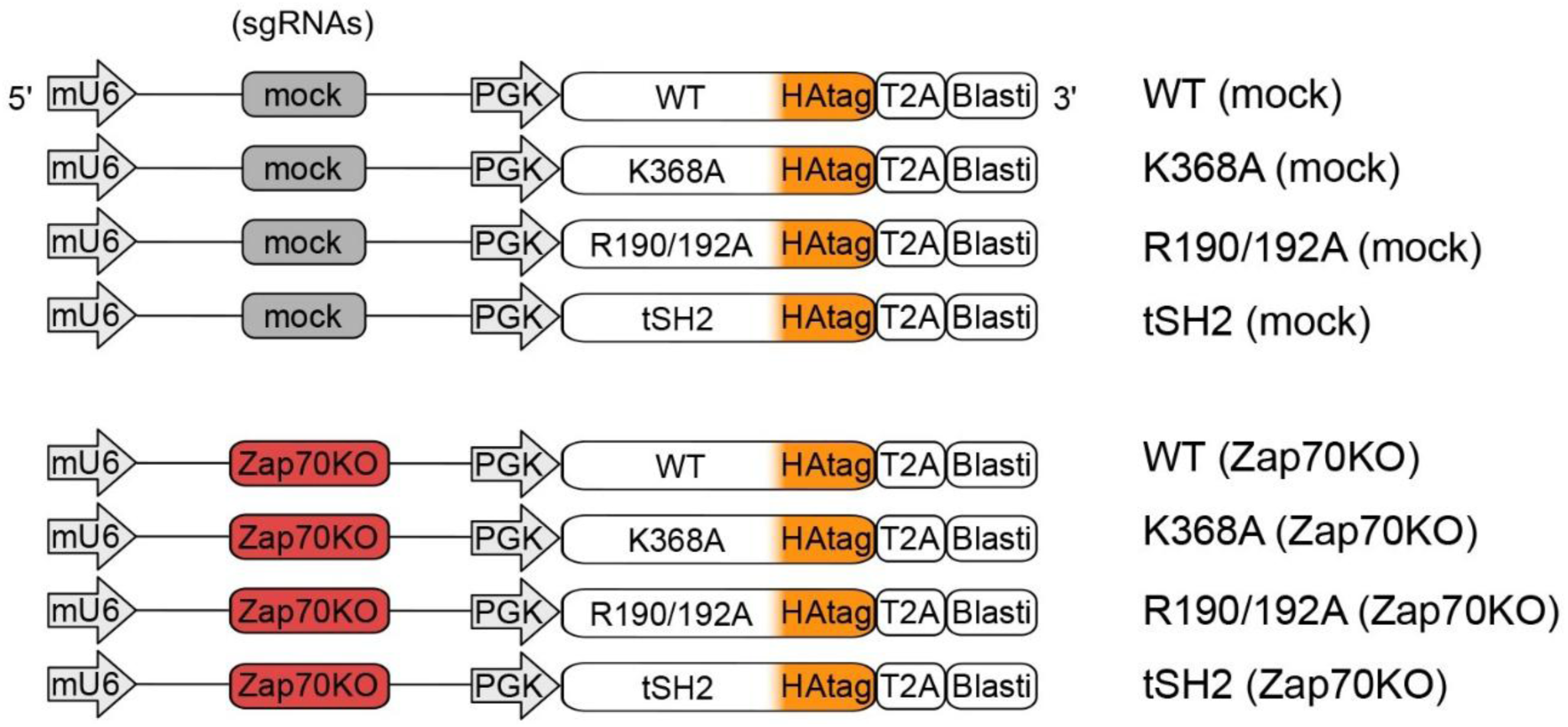
Schematic overview of the Zap70 constructs. Schematic representation of the retroviral constructs co-expressing either a mock or ZAP70-targeting gRNA (Zap70KO), and a PGK promoter-driven HA-tagged ZAP70 variants (WT, K368A, R190/192A, or tSH2), followed by a T2A-linked blasticidin resistance cassette.

**Supplementary Figure 4.**
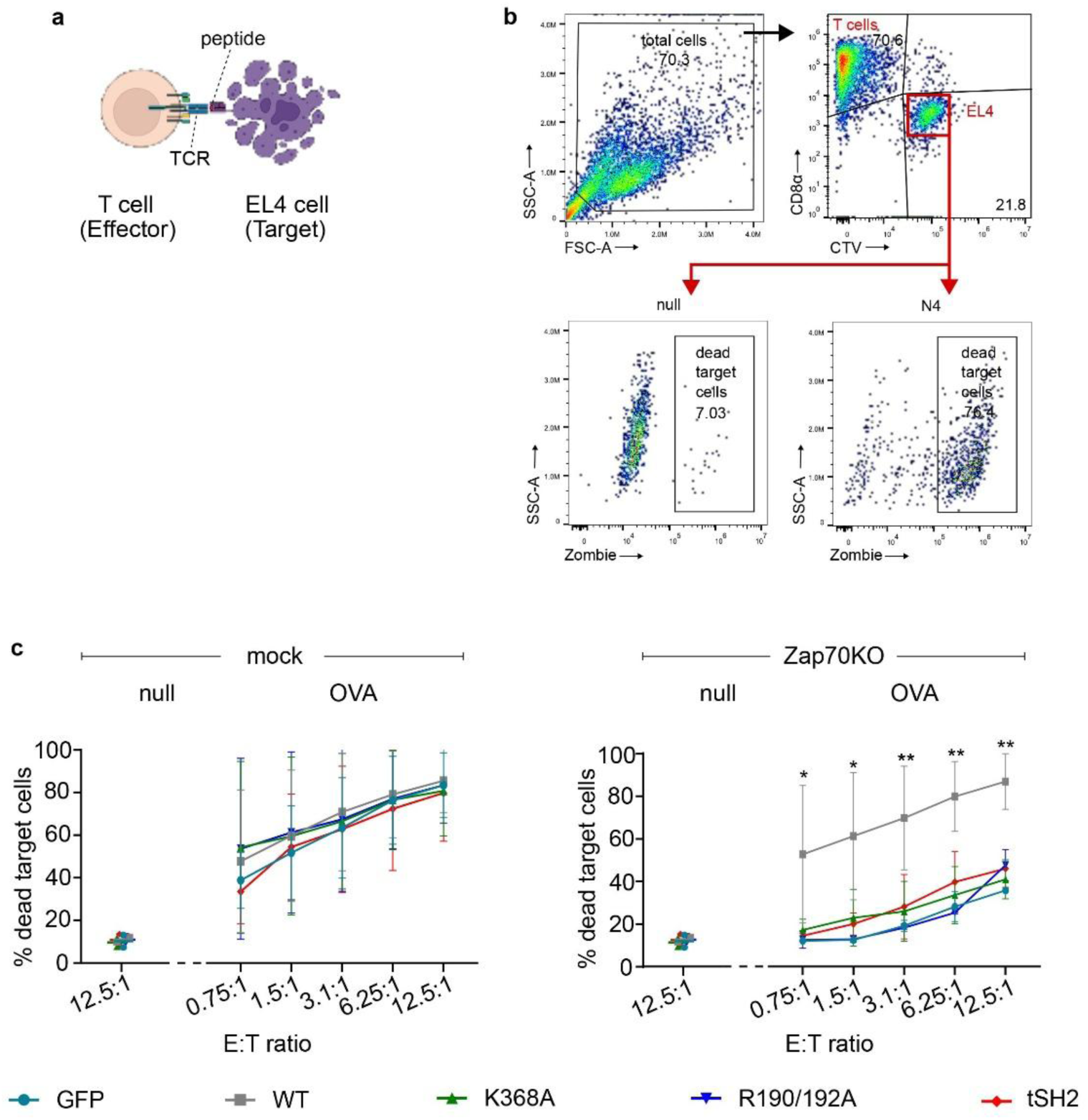
Cytotoxic phenotype of Zap70 mutants requires knockout of endogenous Zap70. OT-I Cas9 T cells were transduced with construct encoding mock gRNA (mock) or gRNA targeting Zap70 (Zap70KO) together with the indicated proteins (GFP, Zap70 WT or Zap70 mutants). Cell Trace Violet-labelled EL4 target cells were loaded with 1μM OVA null peptide, and incubated with transduced T cells at indicated E:T ratios for 4h. (a) Schematic overview of the killing assay. (b) Flow cytometry gating strategy used to analyze target cell death. (c) Percentage of dead target cells after co-culture at indicated E:T ratios of mock (left) or Zap70KO T cells (right), expressing the indicated Zap70 constructs. Data from one experiment, with 3 replicates per condition. Data are presented as mean ± SD. Statistical analysis was performed using two-way ANOVA with Tukey’s multiple comparisons test. P < 0.05 (*), P < 0.01 (**).

**Supplementary Figure 5.**
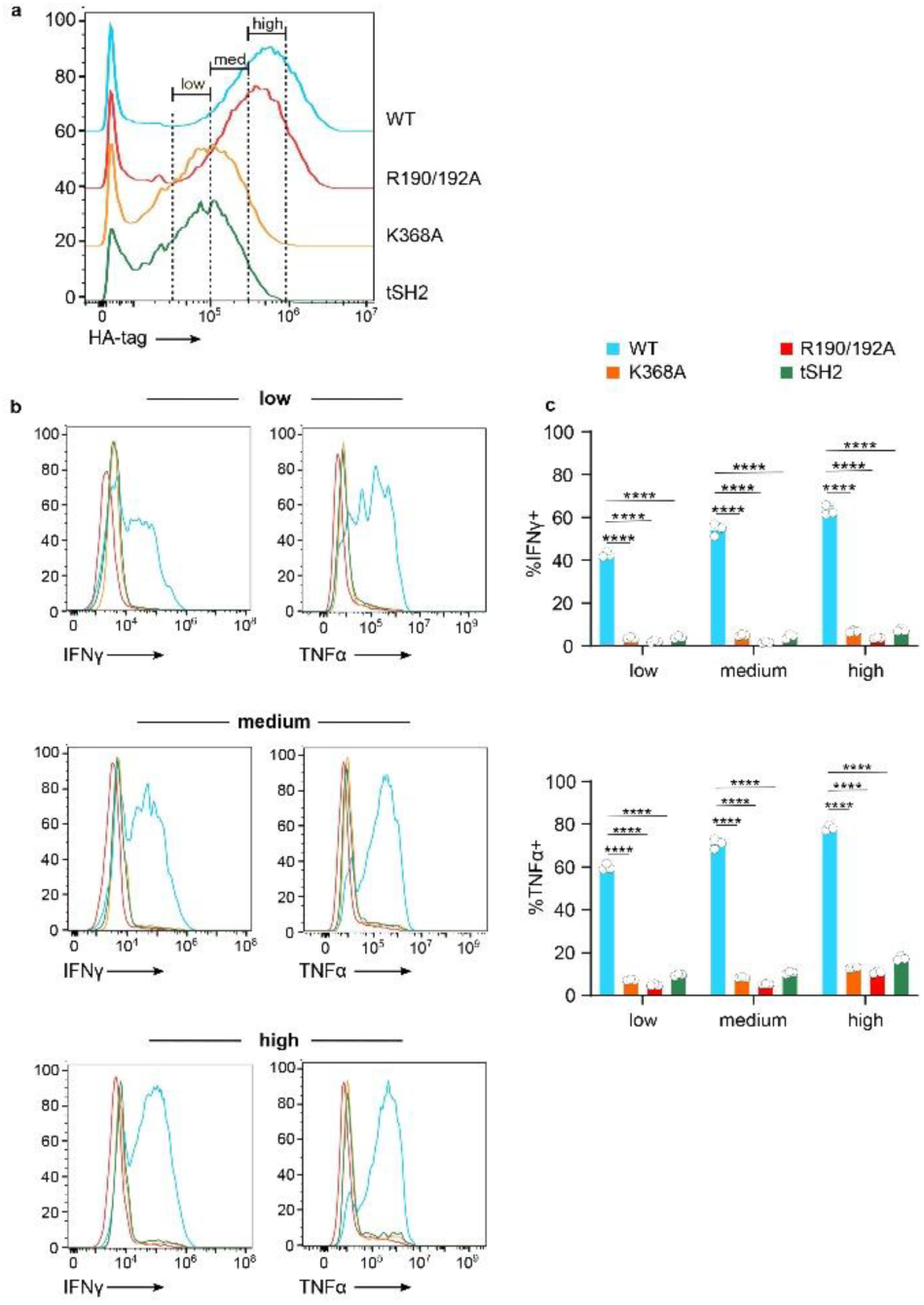
The phenotype of Zap70 mutants is not due to their reduced expression levels. OT-I Cas9 T cells were transduced with construct encoding gRNA targeting Zap70 (Zap70KO) together with the indicated Zap70 constructs. EL4 target cells were loaded with 1nM OVA peptide and used to stimulate transduced T cells. Intracellular staining was used to quantify HA, IFN-γ and TNF-α. (a) Representative flow cytometry histograms showing Zap70low, Zap70med and Zap70high gates. (b) Flow cytometry histograms of IFNγ (left) and TNFα (right) secretion in stimulated T cells expressing low (top), medium (middle) and high (bottom) levels of WT and mutant Zap70. (c) Quantification of T cell percentage expressing IFNγ (top) and TNFα (bottom) in different T cell populations expressing WT and mutant Zap70. Data from one experiment, with 3 replicates per condition. Data are presented as mean ± SD, with each dot representing a replicate. Statistical analysis was performed using two-way ANOVA with Tukey’s multiple comparisons test. P < 0.0001 (****).

## Notes

### Competing Interest Statement

The authors have declared no competing interest.

## References

1. Huang, B., Johansen, K. H. & Schwartzberg, P. L. Efficient CRISPR/Cas9-Mediated Mutagenesis in Primary Murine T Lymphocytes. Curr. Protoc. Immunol. 124, e62 (2019).

2. Oh, S. A., Seki, A. & Rutz, S. Ribonucleoprotein Transfection for CRISPR/Cas9-Mediated Gene Knockout in Primary T Cells. Curr. Protoc. Immunol. 124, e69 (2019).

3. Laprie-Sentenac, M., Cretet-Rodeschini, C. & Menger, L. Optimized protocol to generate genome-wide inactivated Cas9-expressing murine T cells. STAR Protoc. 4, 101922 (2023).

4. Kurachi, M. et al. Optimized retroviral transduction of mouse T cells for in vivo assessment of gene function. Nat. Protoc. 12, 1980–1998 (2017).

5. Vincent, R. L., Li, F., Ballister, E. R., Arpaia, N. & Danino, T. Efficient Generation of Murine Chimeric Antigen Receptor (CAR)-T Cells. (2024). doi:10.3791/65887.

6. Katz, Z. B., Novotná, L., Blount, A. & Lillemeier, B. F. A cycle of Zap70 kinase activation and release from the TCR amplifies and disperses antigenic stimuli. Nat. Immunol. 18, 86–95 (2017).

7. Meana, C., San-José, G., Balboa, M. A. & Casas, J. Spatial regulation of Lck activation at the CD8 immune synapse revealed by a FRET-Based biosensor. Cell. Mol. Life Sci. CMLS 83, (2026).

8. Taylor, J. et al. Exploration of T cell immune responses by expression of a dominant-negative SHP1 and SHP2. Front. Immunol. 14, 1119350 (2023).

9. Johansen, K. H. How CRISPR/Cas9 Gene Editing Is Revolutionizing T Cell Research. DNA Cell Biol. 41, 53–57 (2022).

10. Seki, A. & Rutz, S. Optimized RNP transfection for highly efficient CRISPR/Cas9-mediated gene knockout in primary T cells. J. Exp. Med. 215, 985–997 (2018).

11. Nüssing, S., et al. Efficient CRISPR/Cas9 Gene Editing in Uncultured Naive Mouse T Cells for In Vivo Studies. J. Immunol. Baltim. Md 1950 204, 2308–2315 (2020).

12. Pfenninger, P., Yerly, L. & Abe, J. Naïve Primary Mouse CD8(+) T Cells Retain In Vivo Immune Responsiveness After Electroporation-Based CRISPR/Cas9 Genetic Engineering. Front. Immunol. 13, 777113 (2022).

13. Kornete, M., Marone, R. & Jeker, L. T. Highly Efficient and Versatile Plasmid-Based Gene Editing in Primary T Cells. J. Immunol. Baltim. Md 1950 200, 2489–2501 (2018).

14. Maruyama, T. et al. Increasing the efficiency of precise genome editing with CRISPR-Cas9 by inhibition of nonhomologous end joining. Nat. Biotechnol. 33, 538–542 (2015).

15. Leal, A. F., Herreno-Pachón, A. M., Benincore-Flórez, E., Karunathilaka, A. & Tomatsu, S. Current Strategies for Increasing Knock-In Efficiency in CRISPR/Cas9-Based Approaches. Int. J. Mol. Sci. 25, (2024).

16. Nyberg, W. A. et al. An evolved AAV variant enables efficient genetic engineering of murine T cells. Cell 186, 446–460.e19 (2023).

17. de Menezes, M. N. et al. High efficiency CRISPR knock-in demonstrates that TCF1 is insufficient to reverse T cell exhaustion. Nat. Commun. 17, (2026).

18. Kosaka, M. et al. Evaluation of the loading capacity and patterns of packaged DNA in AAV genomes of different sizes using long-read sequencing. Mol. Ther. Methods Clin. Dev. 33, 101474 (2025).

19. Bing, S. et al. Integrated computational and experimental immunoengineering of adeno-associated virus capsid T cell epitopes in mice. Nat. Commun. 17, (2026).

20. Fujita, T. & Fujii, H. Identification of proteins associated with an IFNγ-responsive promoter by a retroviral expression system for enChIP using CRISPR. PloS One 9, e103084 (2014).

21. Ran, F. A. et al. Genome engineering using the CRISPR-Cas9 system. Nat. Protoc. 8, 2281–2308 (2013).

22. Labun, K. et al. CHOPCHOP v3: expanding the CRISPR web toolbox beyond genome editing. Nucleic Acids Res. 47, W171–W174 (2019).

23. Gascoigne, N. R. J., Casas, J., Brzostek, J. & Rybakin, V. Initiation of TCR phosphorylation and signal transduction. Front. Immunol. 2, 72 (2011).

24. Klammt, C. et al. T cell receptor dwell times control the kinase activity of Zap70. Nat. Immunol. 16, 961–969 (2015).

25. Au-Yeung, B. B., Shah, N. H., Shen, L. & Weiss, A. ZAP-70 in Signaling, Biology, and Disease. Annu. Rev. Immunol. 36, 127–156 (2018).

26. Xie, K., Minkenberg, B. & Yang, Y. Boosting CRISPR/Cas9 multiplex editing capability with the endogenous tRNA-processing system. Proc. Natl. Acad. Sci. U. S. A. 112, 3570–3575 (2015).

27. Dong, F., Xie, K., Chen, Y., Yang, Y. & Mao, Y. Polycistronic tRNA and CRISPR guide-RNA enables highly efficient multiplexed genome engineering in human cells. Biochem. Biophys. Res. Commun. 482, 889–895 (2017).

28. Liu, Z. et al. Systematic comparison of 2A peptides for cloning multi-genes in a polycistronic vector. Sci. Rep. 7, 2193 (2017).

29. Garcillán, B. et al. CD3G or CD3D Knockdown in Mature, but Not Immature, T Lymphocytes Similarly Cripples the Human TCRαβ Complex. Front. Cell Dev. Biol. 9, 608490 (2021).

30. Garcillán, B. et al. The role of the different CD3γ domains in TCR expression and signaling. Front. Immunol. 13, 978658 (2022).

31. Bunnell, S. C. et al. T cell receptor ligation induces the formation of dynamically regulated signaling assemblies. J. Cell Biol. 158, 1263–1275 (2002).

32. Klammt, C. & Lillemeier, B. F. How membrane structures control T cell signaling. Front. Immunol. 3, 291 (2012).

33. Lin, J. J. et al. Membrane Association Transforms an Inert Anti-TCRβ Fab’ Ligand into a Potent T Cell Receptor Agonist. Biophys. J. 118, 2879–2893 (2020).

34. Qian, D., Mollenauer, M. N. & Weiss, A. Dominant-negative zeta-associated protein 70 inhibits T cell antigen receptor signaling. J. Exp. Med. 183, 611–620 (1996).

35. Park, M.-J. et al. SH2 Domains Serve as Lipid-Binding Modules for pTyr-Signaling Proteins. Mol. Cell 62, 7–20 (2016).

36. Juang, J. et al. Peptide-MHC heterodimers show that thymic positive selection requires a more restricted set of self-peptides than negative selection. J. Exp. Med. 207, 1223–1234 (2010).

37. Kotov, D. I. et al. TCR Affinity Biases Th Cell Differentiation by Regulating CD25, Eef1e1, and Gbp2. J. Immunol. Baltim. Md 1950 202, 2535–2545 (2019).

38. Kotov, J. A. et al. BCL6 corepressor contributes to Th17 cell formation by inhibiting Th17 fate suppressors. J. Exp. Med. 216, 1450–1464 (2019).

39. Dubrot, J. et al. In vivo screens using a selective CRISPR antigen removal lentiviral vector system reveal immune dependencies in renal cell carcinoma. Immunity 54, 571–585.e6 (2021).

40. Bresser, K. et al. A mouse model that is immunologically tolerant to reporter and modifier proteins. Commun. Biol. 3, 273 (2020).

41. Ewaisha, R. & Anderson, K. S. Immunogenicity of CRISPR therapeutics-Critical considerations for clinical translation. Front. Bioeng. Biotechnol. 11, 1138596 (2023).

42. Friedman, R. S., Beemiller, P., Sorensen, C. M., Jacobelli, J. & Krummel, M. F. Real-time analysis of T cell receptors in naive cells in vitro and in vivo reveals flexibility in synapse and signaling dynamics. J. Exp. Med. 207, 2733–2749 (2010).

43. Roncagalli, R. et al. Quantitative proteomics analysis of signalosome dynamics in primary T cells identifies the surface receptor CD6 as a Lat adaptor-independent TCR signaling hub. Nat. Immunol. 15, 384–392 (2014).

44. Shen, L., Matloubian, M., Kadlecek, T. A. & Weiss, A. A disease-associated mutation that weakens ZAP70 autoinhibition enhances responses to weak and self-ligands. Sci. Signal. 14, (2021).

45. Dong, M. B. et al. Systematic Immunotherapy Target Discovery Using Genome-Scale In Vivo CRISPR Screens in CD8 T Cells. Cell 178, 1189–1204.e23 (2019).

46. LaFleur, M. W. et al. PTPN2 regulates the generation of exhausted CD8(+) T cell subpopulations and restrains tumor immunity. Nat. Immunol. 20, 1335–1347 (2019).

47. Garcia, J. et al. Naturally occurring T cell mutations enhance engineered T cell therapies. Nature 626, 626–634 (2024).

48. Orozco, R. C., Marquardt, K., Mowen, K. & Sherman, L. A. Proautoimmune Allele of Tyrosine Phosphatase, PTPN22, Enhances Tumor Immunity. J. Immunol. Baltim. Md 1950 207, 1662–1671 (2021).

49. LaFleur, M. W. et al. In Vivo CRISPR Screening Reveals CHD7 as a Positive Regulator of Short-lived Effector Cells. J. Immunol. Baltim. Md 1950 213, 1528–1541 (2024).

